# Fast-slow traits predict competition network structure and its response to resources and enemies

**DOI:** 10.1101/2023.11.07.565954

**Authors:** Caroline Daniel, Eric Allan, Hugo Saiz, Oscar Godoy

## Abstract

Plants interact in complex networks but how their structure depends on resources, natural enemies and species resource-use strategy remains poorly understood. Here, we quantified competition networks among 18 plants varying in fast-slow strategy, by testing how increased nutrient availability and reduced foliar pathogens affected intra- and inter-specific interactions. Our results show that nitrogen and pathogens altered several aspects of network structure, often in unexpected ways due to fast and slow growing species responding differently. Nitrogen addition increased competition asymmetry in slow growing networks, as expected, but decreased it in fast growing networks. Pathogen reduction made networks more even and less skewed because pathogens targeted weaker competitors. Surprisingly, pathogens and nitrogen dampened each other’s effect. Our results show that plant growth strategy is key to understand how competition respond to resources and enemies, a prediction from classic theories which has rarely been tested by linking functional traits to competition networks.

## Introduction

Global changes are dramatically altering ecological communities (Brooker 2006). These changes can occur through shifts in plant performance (Ahmad, Diwan, and Abrol 2010) and alterations in plant-plant interactions (Matías et al. 2018; Van Dyke, Levine, and Kraft 2022). While past research has extensively investigated how resource gradients (e.g., Tilman 1985; DiTommaso and Aarssen 1989) or plant consumers can change plant interactions (e.g., Holt, Grover, and Tilman 1994), most studies have either quantified overall competitive responses of individual species (Dormann and Roxburgh 2005; Yang et al. 2022), or have examined pairwise interactions among a limited number of species (Chesson 2000). Consequently, our understanding of how species interactions change within communities of multiple interacting species remains limited (Levine et al. 2017). This knowledge is essential to gain a more mechanistic understanding of global change effects on the assembly, stability and functioning of ecological communities (Loreau and Hector 2001; Mayfield and Levine 2010; López-Angulo et al. 2018).

Initial efforts to upscale the study of pairwise interactions to multispecies plant-plant networks were largely theoretical (Allesina and Levine 2011). Empirical examples focused on quantifying some aspects of the competition network (Gallien et al. 2017; Soliveres et al. 2015; Saiz et al. 2019; Kinlock 2019) but did not look at how multiple metrics of network structure vary with environmental conditions. To characterise interaction networks in contrasting conditions (Table 1) we need to incorporate two key aspects of plant-plant interactions. First, we need to simultaneously account for negative and positive interactions, as both are prevalent in natural ecosystems (Losapio et al. 2021; Bimler et al. 2018). Second, we need to quantify plant-plant interactions as per-capita effects to assess their demographic consequences (abundance shifts, local extinction events, etc.). Both aspects are crucial to understand the long-term consequences of changes in plant-plant interactions.

**Table 1.**
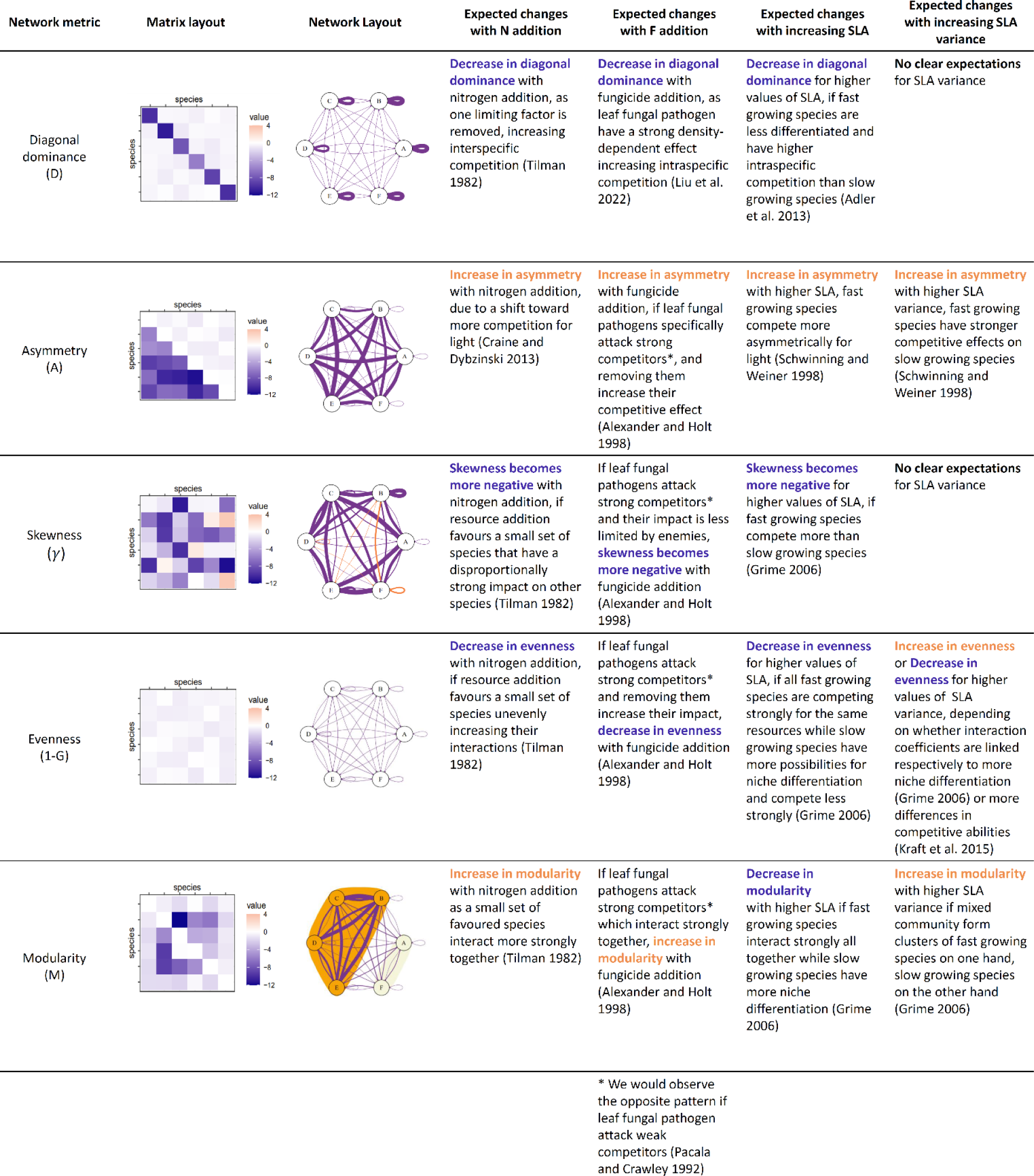
Network metrics selected for the study and their corresponding hypotheses for how they will respond to treatments and strategies (N = nitrogen, F = Fungicide, SLA = Specific Leaf Area). Every metric is presented with an example of the competition matrix and the typical network layout when the metric is high (i.e., the network is highly diagonal dominant, highly asymmetric, highly even, highly skewed, highly modular). Example matrices and networks are presented for a species richness of n = 6 species, for better clarity.

| Network metric | Matrix layout | Network Layout | Expected changes with N addition | Expected changes with F addition | Expected changes with increasing SLA | Expected changes with increasing SLA variance |
| --- | --- | --- | --- | --- | --- | --- |
| Diagonal dominance (D) |  |  | Decrease in diagonal dominance with nitrogen addition, as one limiting factor is removed, increasing interspecific competition (Tilman 1982) | Decrease in diagonal dominance with fungicide addition, as leaf fungal pathogens have a strong density-dependent effect increasing intraspecific competition (Liu et al. 2022) | Decrease in diagonal dominance for higher values of SLA, if fast growing species are less differentiated and have higher intraspecific competition than slow growing species (Adler et al. 2013) | No clear expectations for SLA variance |
| Asymmetry (A) |  |  | Increase in asymmetry with nitrogen addition, due to a shift toward more competition for light (Craine and Dybzinski 2013) | Increase in asymmetry with fungicide addition, if leaf fungal pathogens specifically attack strong competitors*, and removing them increase their competitive effect (Alexander and Holt 1998) | Increase in asymmetry with higher SLA, fast growing species compete more asymmetrically for light (Schwinning and Weiner 1998) | Increase in asymmetry with higher SLA variance, fast growing species have stronger competitive effects on slow growing species (Schwinning and Weiner 1998) |
| Skewness ($\gamma$) | | | Skewness becomes more negative with nitrogen addition, if resource addition favours a small set of species that have a disproportionately strong impact on other species (Tilman 1982) | If leaf fungal pathogens attack strong competitors* and their impact is less limited by enemies, skewness becomes more negative with fungicide addition (Alexander and Holt 1998) | Skewness becomes more negative for higher values of SLA, if fast growing species compete more than slow growing species (Grime 2006) | No clear expectations for SLA variance |
| Evenness (1-G) |  |  | Decrease in evenness with nitrogen addition, if resource addition favours a small set of species unevenly increasing their interactions (Tilman 1982) | If leaf fungal pathogens attack strong competitors* and removing them increase their impact, decrease in evenness with fungicide addition (Alexander and Holt 1998) | Decrease in evenness for higher values of SLA, if all fast growing species are competing strongly for the same resources while slow growing species have more possibilities for niche differentiation and compete less strongly (Grime 2006) | Increase in evenness or Decrease in evenness for higher values of SLA variance, depending on whether interaction coefficients are linked respectively to more niche differentiation (Grime 2006) or more differences in competitive abilities (Kraft et al. 2015) |
| Modularity (M) |  |  | Increase in modularity with nitrogen addition as a small set of favoured species interact more strongly together (Tilman 1982) | If leaf fungal pathogens attack strong competitors* which interact strongly together, increase in modularity with fungicide addition (Alexander and Holt 1998) | Decrease in modularity with higher SLA if fast growing species interact strongly all together while slow growing species have more niche differentiation (Grime 2006) | Increase in modularity with higher SLA variance if mixed community form clusters of fast growing species on one hand, slow growing species on the other hand (Grime 2006) |

\* We would observe the opposite pattern if leaf fungal pathogens attack weak competitors (Pacala and Crawley 1992)

Nitrogen addition is a key global change driver that can dramatically change the overall structure of plant-plant interactions. A large body of research has shown that nitrogen addition decreases coexistence opportunities by removing a limiting factor for plant growth (Tilman 1982) and leads to long-term diversity loss (Suding et al. 2005; Isbell et al. 2013; Crawley et al. 2005). Nitrogen could therefore impact network structure in several ways: removing a limiting factor will reduce intraspecific interactions relative to interspecific ones (Adler et al. 2018), leading to reduced diagonal dominance in the competition matrix. However, nitrogen addition alone might not strongly reduce niche dimensionality if other resources remain limiting (Harpole et al. 2016). In addition, nitrogen addition often makes plants taller leading to an increase in light competition (Hautier, Niklaus, and Hector 2009; Eskelinen et al. 2022). This is likely to increase the asymmetry of interactions by promoting competitive superiority for taller species (DeMalach, Zaady, and Kadmon 2017). Overall, nitrogen addition is expected to impact several aspects of plant interaction networks (see Table 1 for detailed information) and we lack empirical data on how several metrics respond.

Consumers, such as herbivores or pathogens, can also alter plant-plant interactions (Chesson and Kuang 2008). The existence of shared consumers can lead to apparent competition among plant species (Holt 1977) with potentially opposing effects on plant-plant interaction networks. On the one hand, plant enemies, like specialist fungal pathogens, can maintain biodiversity by increasing intraspecific relative to interspecific competition (Janzen 1970; Connell 1971; Bagchi et al. 2014), thereby increasing diagonal dominance. Pathogens might also reduce the growth of the most competitive species (Alexander and Holt 1998) causing plant-plant interactions to become weaker, less negatively skewed, and more evenly distributed. On the other hand, pathogens could reduce biodiversity by decreasing the ability of weaker competitors to tolerate competition (Pacala and Crawley 1992; Mordecai 2011; Parker and Gilbert 2018). This would result in an increase in the negative skewness of the network, meaning more extreme competition coefficients, and a decrease in the evenness of plant-plant interactions. Because studies on foliar fungal pathogens have found inconsistent effects on plant communities (Spear and Mordecai 2018; Allan, van Ruijven, and Crawley 2010; Liu et al. 2022; Granjel, Allan, and Godoy 2023) both alternatives seem plausible. Finally, fungal pathogens might interact with resources (Allan and Crawley 2011; Cleland and Harpole 2010), and they resources and pathogens might amplify or dampen each other’s effects, depending on whether they favour the same set of species or not. Only experiments crossing nitrogen and enemy removal can address these potential interactions.

Plant-plant interactions might also vary between plants with different resource-use strategies. A key axis of plant functional variation separates fast growing, resource acquisitive species from slow growing, conservative species with high levels of defence against consumers (Coley, Bryant, and Chapin 1985; Poorter, Remkes, and Lambers 1990). Fast growing species can rapidly acquire resources and are strongly competitive for light, whereas slow growing species are adapted to low resource conditions and may be more competitive for soil resources. Several traits relate to the resource economics spectrum and specific leaf area (SLA) is one of the most commonly measured (Kunstler et al. 2016; Funk et al. 2017; Díaz et al. 2016). Both the variation in SLA between the plants in a network (community) and the average SLA of the plants in the network could determine its structure. First, if competitive ability depends on SLA (Kraft, Godoy, and Levine 2015), we should expect high evenness between species varying minimally in SLA and more asymmetric competition in networks with species varying strongly in SLA (Van Dyke, Levine, and Kraft 2022). In addition, modularity might be higher in networks with high SLA variation, since we species with similar SLA should form cluster with similar resource use (Gross et al. 2009). Second, networks of fast-growing plants (high mean SLA) competing for light might be more asymmetrical and skewed (DeMalach, Zaady, and Kadmon 2017), whereas networks of slow growing species that differ in nutrient niches might be more even with higher diagonal dominance (Tilman 1982). As fast and slow species are expected to respond differently to pulses of resources and to enemy attack (Loranger et al. 2012; Grigulis et al. 2013), traits linked to the resource economics spectrum could also predict the response of competition networks to resources and enemies. For instance, fast species are likely to benefit more from nutrient addition (da Silveira Pontes et al. 2010) and a reduction in enemies (Coley, Bryant, and Chapin 1985; Cappelli et al. 2020). Many studies have compared interactions between high and low resource environments, but they have not distinguished whether plant-plant interactions change because overall resources levels (nitrogen) changes or because functional composition changes (from slow to fast species).

Here we investigate how nitrogen enrichment and the foliar fungal pathogen removal (with fungicide) alter interaction networks between plants differing in their resource economics strategy. Networks were built from pairwise intra- and inter-specific per capita effects among 18 perennial plant species, differing strongly in SLA, measured in a large grassland experiment (PaNDiv Experiment, Switzerland). We quantified 18×18 species competition networks in the four combinations of nitrogen and fungicide treatments, in late spring and late summer. Environmental conditions are highly variable throughout the season, so we tested how consistent the patterns are across seasons varying in climatic conditions, especially in water availability. We therefore investigated: (1) what attributes of the whole plant network (Table 1) varied most with nitrogen addition and pathogen reduction, alone or in combination; (2) whether interaction networks composed of either slow growing, fast growing or a mix of fast and slow growing species (i.e., differing in the variance and mean of SLA) responded differently to nitrogen and foliar pathogens.

## Material and Methods

### Experimental set-up

We conducted our study within the PaNDiv experiment in Bern, Switzerland. PaNDiv experiment was set up in October 2015 and contains 336 plots of 2m × 2m with different numbers and compositions of 20 perennial grass and herb species, selected to vary in their SLA and leaf nitrogen content and therefore in resource use strategy (Table S1). In total, 80 monoculture plots and 256 mixture plots were established, varying in functional composition and species richness (for more information, see Pichon et al. 2020). Species richness and functional composition are crossed with four other treatments: control (C), nitrogen addition (N), fungicide addition (F) and combined nitrogen and fungicide addition (NF). Fertilised plots received N in the form of urea twice a year in April and late June, for an annual addition of 100 kg N ha^-1^y^-1^. Two fungicides (“Score Profi”, 23.5 % Difenoconazol 250 g.L^-1^ and “Heritage Flow”, 22.8% Azoxystrobin 400 g.L^-1^) were sprayed four times a year (early April, early June, late July and September) to reduce foliar pathogens. Water was sprayed simultaneously on the untreated plots. The experiment was weeded three times a year to maintain the diversity and composition treatments. Plots were mown twice a year, in mid-June and mid-August, which corresponds to intermediate to extensive grassland management (Blüthgen et al. 2012).

### Phytometer experiment to estimate species interactions

We took advantage of this experimental design to estimate species interactions as pairwise per-capita intra and interspecific effects. We used a phytometer approach, which involved measuring biomass production of individuals planted in neighbourhoods differing in density (from no neighbours to crowded neighbourhoods) and relative frequency of different species (from growing only with conspecific neighbours to heterospecific ones), across the four different treatments. The per-capita effects indicate competition (or facilitation) that occurs when the phytometer biomass of a given species *i* is reduced (or increased) as the neighbourhood density of species *j* increases. Neighbourhoods were defined as all plants within a 20 cm radius of the phytometer (Granjel, Allan, and Godoy 2023). We obtained phytometers by germinating commercially supplied seeds (UFA Samen in Switzerland and Rieger Hoffmann in Germany) in an experimental greenhouse (Ostermundigen, Switzerland August 2019). We planted phytometers in the field in September 2019 when they had grown to 3-5 cm tall. Two of the 20 species, *Anthriscus sylvestris* and *Heracleum sphondylium*, did not germinate well enough, and could not be included. However, these species also did not establish well on the PaNDiv experiment (Cappelli et al. 2022). The remaining 18 species were each planted into 10 different intraspecific neighbourhoods varying in density, using monoculture plots, and into three different neighbourhoods for each heterospecific competitor. Three species, *Poa trivialis*, *Anthoxanthum odoratum* and *Rumex acetosa*, did not occur in high density neighbourhoods, although they were present in at least some neighbourhoods for each phytometer. Finally, we grew phytometers of all species in plots with no neighbours, to estimate growth without any other individuals present. Phytometers were planted 20 cm apart in 2×2m plots covered with landscape fabric to prevent weed growth and received the four combinations of nitrogen and fungicide (as applied on PaNDiv). They were cut at the same time as the field was mown. Overall, we planted a total of 4’248 phytometers for all species, which corresponded to 3’240 heterospecific neighbourhoods (3 replicates × 4 treatments × 18 species × 15 neighbourhoods), 720 conspecific neighbourhoods (10 replicates × 4 treatments × 18 species) and 288 no competition neighbourhoods (4 replicates × 4 treatments × 18 species). These phytometers were measured in June 2020 two-three weeks before the mowing, and again in August 2020 after regrowth. We started with 4’248 phytometers planted in September 2019, and recovered 3’888 (91.5%) in June 2020 and 2’722 (64.1%) in August 2020.

### Statistical approach to estimate species interactions

In order to obtain the matrix of species interactions and assess per capita effects between pairs of species, we related phytometer biomass to the density of species present within its neighbourhood. For each of the 18 focal species, we fitted a model with a negative binomial function (‘GLMMTMB’ package, version 1.1.2.3, Brooks et al. 2017). This function was selected because it allows us to quantify with equal probability both competition and facilitation. In our analyses, the response variable (phytometer biomass as dry weight in mg.) was related to (1) cover of all neighbour species (including conspecifics), (2) treatment (control, nitrogen, fungicide and combined nitrogen and fungicide) and (3) sampling time (June or August). We also include as (4) cover of weeds, i.e., plants other than the 18 target species, in the neighbourhoods as a covariate, however, weed cover was generally low (5.39% on average). The model also included all interaction effects between treatment, sampling period and neighbour species, except for the interaction between sampling period and treatment, which was not estimated due to convergence issues. The level of experimental replication was sufficient to independently estimate each interspecific pairwise interactions from neighbourhoods in which several competitors were co-occurring, however some interspecific coefficients, particularly those between species that were rare in the experiment, were estimated with more error. To account for this uncertainty, we divided each interaction coefficient by its standard error, which meant that poorly estimated coefficients became close to 0. We therefore estimated all intraspecific effects (α_*ii*_, the per capita effect of one species on itself) and pairwise interspecific effects (α_*ij*_, the per capita effect of species j on species i) across treatments and sampling times. In total, we built eight matrices (4 treatments × 2 sampling times) of 18 × 18 species, with 4 × 2 × 18^2^ = 2’592 interaction coefficients. These coefficients were used to compute several network metrics (Table 1). All analyses were done using R version 4.1.0.

### Computation of network metrics

We selected five different metrics that represent several features of the plant networks (see detailed explanation in Table 1). Diagonal dominance (*D*) and asymmetry (*A*) were calculated following equations 1 and 2 (Box 1), other metrics were calculated following equations 3-5 (Box 1). Skewness (γ) was calculated with the skewness function of the package ‘moments‘ (version 0.14.1, Komsta and Novomestky 2015). The Gini coefficient (G) was calculated with the function Gini_RSV of the package ‘GiniWegNeg‘ (version 1.0.1, Raffinetti, Siletti, and Vernizzi 2015). Modularity (M) was calculated with the modularity function of the ‘igraph’ package 1.3.1. on the clusters determined with the cluster_optimal function (Csardi 2013), using the absolute values of the coefficients 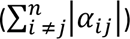 as weights of the edges (Reichardt and Bornholdt 2006).

### Effects of nitrogen, fungicide and species resource use on plant-plant network properties.

We aimed to compare how plant-plant networks varied with nitrogen and fungicide treatments and whether fast and slow growing networks responded differently. To address this, we focused on subnetworks of five species because this is the average number of species found in interspecific neighbourhoods (mean = 5.19 species). We estimated the network metrics described above (Table 1, Box 1) for all 68’916 five species networks composed from the 18 species, in each of the four treatments and two sampling periods. We characterised the growth strategy of these networks using the mean and variance in SLA between the species (Supplementary Method 1). We then fitted models to predict the network metrics for the 55’1328 networks using: (1) nitrogen addition, fungicide addition and their combination (2) the sampling period (June and August), (3) the mean and variance in community SLA and (4) interactions between mean and variance in SLA and the treatments and seasons. The networks of five species are not fully independent since they share species, so we used multi-membership models to correct for the degree of similarity between networks inside the random effects of the mixed model. Multi-membership models are particularly useful when elements which are nested in the random effects are more important than the overall differences between levels of the random factor (here, the nested species identities are likely to be more important than the community combination itself). By considering species identity nested in our communities, we can correct for the various degrees of similarity between networks, especially in the case of one species having a disproportionate effect on the community (Hadfield 2010). We fitted the random effect by following the method described in Ben Bolker’s Github repository (https://github.com/bbolker/mixedmodels-misc/tree/master/notes, “multimember.Rmd” file). We implemented the presence/absence matrix of species in the communities as a random factor in a linear mixed models fitted with the package lme4 version 1.1-27-1 (Bates et al. 2015), and we ran the models and optimised the estimation of the maximum-likelihood coefficients with ‘devTool’ package version 2.4.3 (Wickham et al. 2021).

In the last step of the analyses, we tested whether previous results based on five species would differ if larger networks were considered. Accordingly, we computed all metrics for every possible network containing 5, 7, 11, and 15 species, resulting in a total of n = 292 128 networks. To make this analysis comparable across species richness levels, we focused on networks containing mixed resource use strategies (at least one fast and one slow growing species). This is because networks with 11 or 15 species have to include both strategies, as there are 8 fast and 10 slow growing species, and the range in mean and variance in SLA would therefore be lower in these larger networks. For this last analysis, we also fitted multi-membership models but we only included interactions measured in June.

## Results

We estimated 324 pairwise interaction coefficients for each sampling period and treatment, i.e., 2592 interaction coefficients in total. Overall, 84% (min = 81%, max = 90%) of these interactions were negative (i.e., competitive interactions) and their average strength was -1.55 (min = -10.86, max = 3.00) (Fig. 1 for June and S1 for August). An average competitive interaction of -1.55 corresponds to a reduction of around 26% of the phytometer biomass when competitor cover increases from 0 to 20%. Among all 18 species, *Holcus lanatus* (Hl) had particularly large competitive effects on all others. *Achillea millefolium* (Am) and, to a lesser extent *Holcus lanatus*, were particularly sensitive to competition, while *Rumex acetosa* (Ra) was fairly insensitive to competition.

**Figure 1.**
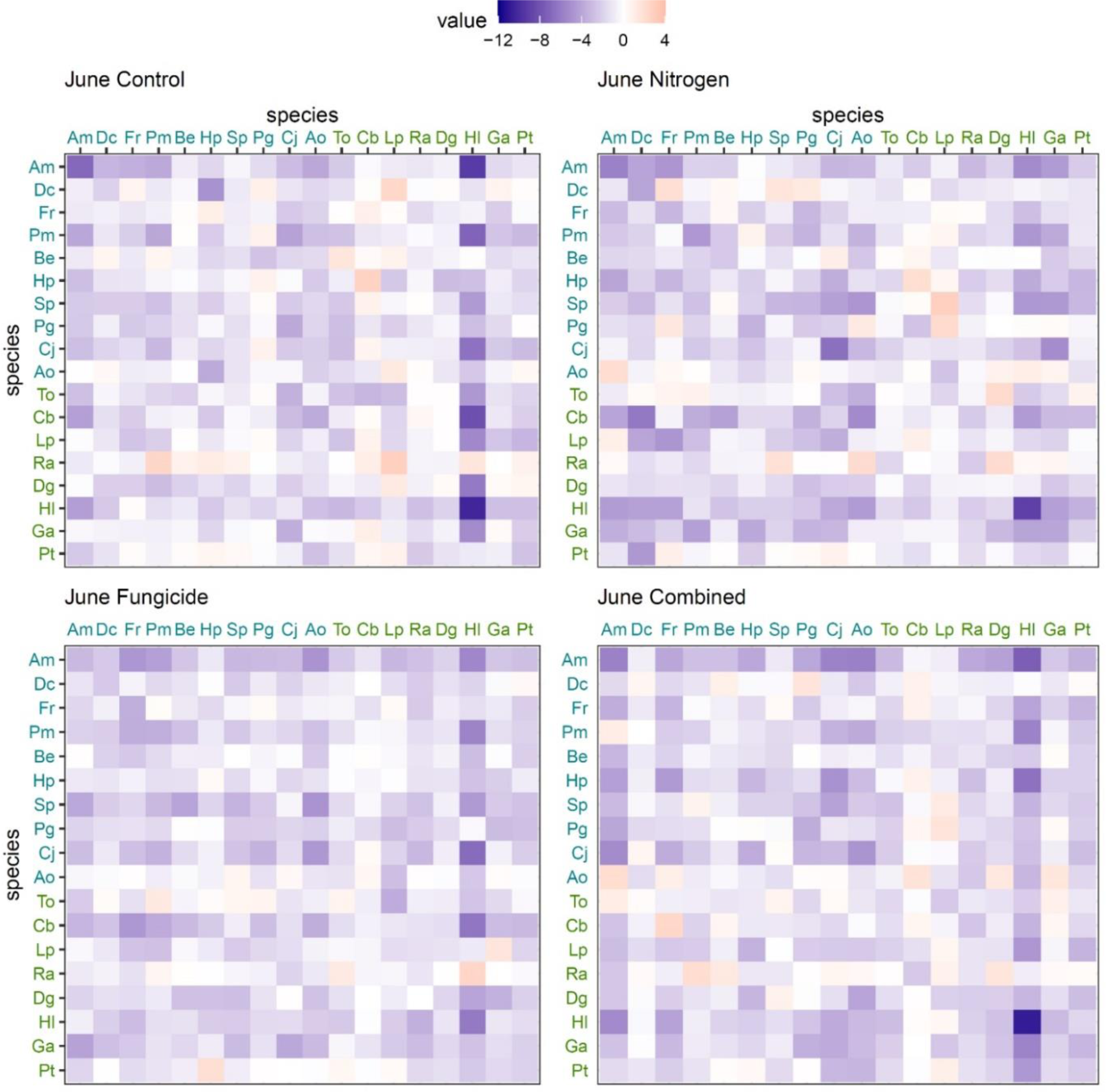
Matrices of pairwise interactions between the 18 species sampled in our experiment in June. August matrices for each treatment are included in Supplementary Figure 1. Each cell shows a pairwise interaction, colours show the sign and the magnitude of the interaction. Species names in blue are slow growing species, green are fast growing species. Columns show species effects on others while rows show species response to competition from others. For instance, row 2 column 1 shows the response of Salvia pratensis to competition from Achillea millefolium and row 1 column 2 shows the effect of Sp on Am. See Supplementary Table S1 for species names abbreviation.

Nitrogen and fungicide addition had significant impacts on almost all network metrics (Figs. 2 and 3, Table S2). As we analysed a very large number of networks, effects were typically significant and we therefore focus only on those effects that we define as “strong”, i.e., with standardised coefficients > 0.1 or < -0.1. Fungicide had the strongest effects and impacted all metrics except asymmetry. Fungicide strongly increased evenness and reduced modularity and negative skewness. Nitrogen addition strongly reduced the diagonal dominance of the networks and increased asymmetry and negative skewness. However, nitrogen also strongly increased evenness and modularity. Nitrogen and fungicide also interacted with each other and often dampened each other’s effects. We analysed whether such dampening was due to different species responding to the two treatments but found no consistent pattern (Figs. S2 and S3). Season primarily affected modularity and networks were strongly modular in August (Figs. S4 and S5).

**Figure 2.**
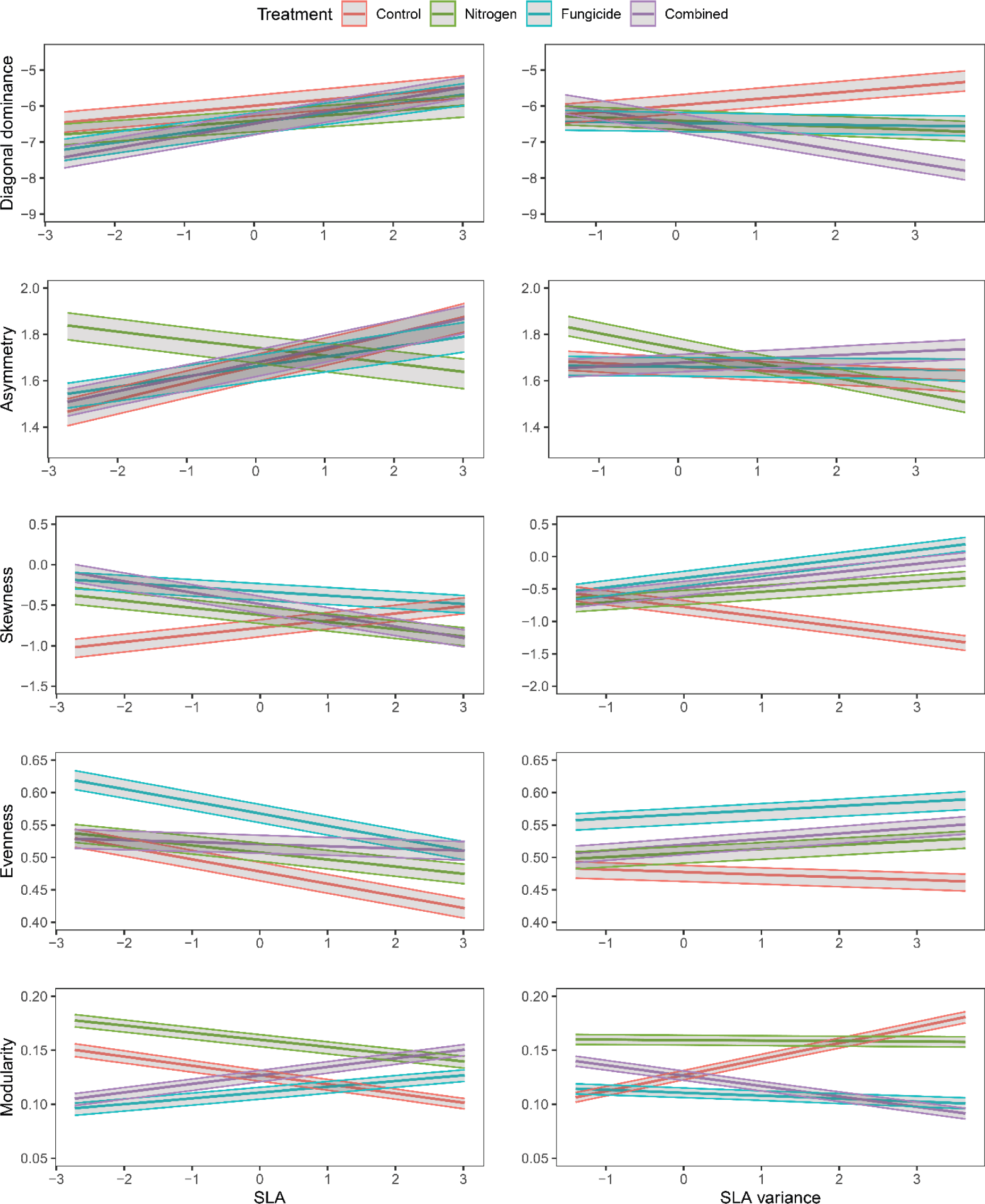
Impact of SLA (left panels), SLA variance (right panels) and treatments on network metrics for communities of five species over the two sampling seasons. For each network metric, colours determine treatments, and model predictions are shown in the form of predicted mean (bold lines) and confidence intervals (upper and lower lines). SLA and SLA variance were scaled for comparison. SLA = specific leaf area.

**Figure 3.**
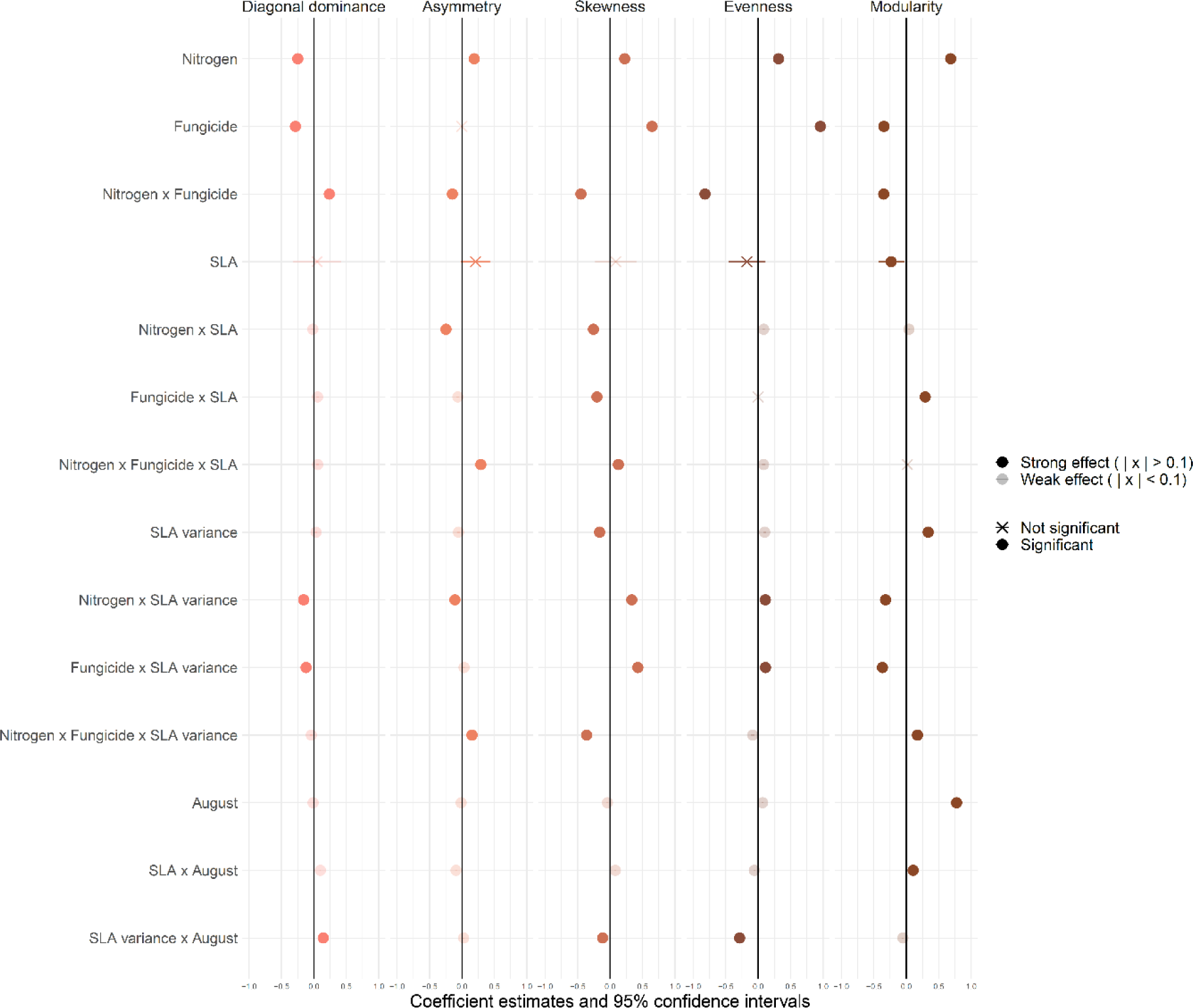
Estimates of fixed effects from multi-membership LMER models showing the effects of nitrogen addition, fungicide addition, SLA, SLA variance and season on functionally mixed communities of 5 species. All network metrics, SLA and SLA variance were scaled for comparison. SLA = specific leaf area.

Communities with different mean SLA often responded in opposing ways to the treatments. Asymmetry increased with nitrogen addition for networks with low SLA, but was reduced with nitrogen in networks with high SLA. Conversely, fungicide reduced modularity in low SLA networks, but led to more modular networks in high SLA communities. A triple interaction was also observed for skewness between average SLA, nitrogen and fungicide addition. This means that skewness was increased by all treatments in low SLA communities but it was reduced, or not affected by the treatments, in high SLA communities. The variance of SLA within communities also strongly affected network metrics response to both nitrogen and fungicide addition (Table S2). Overall, communities with high variance in SLA were more negatively skewed and modular than those with low variance in SLA (Figs. 2 and 3, Table S2). However, both nitrogen and fungicide addition reduced diagonal dominance and increased skewness in communities with high variance in SLA, whereas the treatments had no effect at low variance in SLA. In contrast, evenness was increased by nitrogen and fungicide and modularity was reduced, particularly in communities with high variance in SLA (Fig. 2). Asymmetry was also increased by nitrogen alone, in communities with low variance in SLA, and reduced by nitrogen at high variance in SLA.

Finally, we observed that network metrics changed as we increased the number of species in the network from 5 to 15. However, the treatment effects and interactions between treatments and mean and variance in SLA remained qualitatively the same (Fig. S6 and S7, Table S3).

## Discussion

We found that resource addition and enemy removal strongly impacted plant-plant interaction network structure. Nitrogen addition and the removal of foliar fungal pathogens with fungicide generally reduced the dominance of intraspecific interactions. Nitrogen addition also generally made communities more asymmetric in interspecific competition. Both effects are in line with theoretical expectations that resource limitation and enemies maintain diversity by increasing self-limiting effects and equalising competitive ability between species (Chesson 2000; Buche et al. 2022). However, nitrogen and fungicide also had a range of effects on interaction networks, which did not always agree with major theories (Table 1). We found that both nitrogen and fungicide made communities more even and less negatively skewed, which was counter to our expectations, and they typically dampened each other’s effects, rather than acting in a similar way, as expected. Some of these unexpected results arose because the impacts of nitrogen and fungicide addition were strongly dependent on the mean and variance of specific leaf area (SLA) between species in the networks (Fig. 2). This indicates that overall fast-slow resource strategy is a key driver of plant-plant interactions and determines how they respond to changes in resources and enemies. This idea is implicit in some classic theories (Tilman 1982; Grime 1979) but has rarely been addressed in studies linking functional traits to competition.

Nitrogen addition effects on competition networks strongly depended on the growth strategy of the constituent species. We expected plants to grow taller and compete more for light following nitrogen addition (Eskelinen et al. 2022), which would result in more asymmetric interactions, as tall species strongly affect their neighbours (strong competitive effect) but are only weakly affected by smaller neighbours (weak competitive response). Many studies have looked at traits determining overall competitive effect and response (Goldberg and Landa 1991; Schwinning and Weiner 1998; Wang et al. 2010), and some have identified increased competitive asymmetry as a major mechanism causing species loss following nitrogen enrichment (DeMalach, Zaady, and Kadmon 2017; Xiao et al. 2021), however, few have looked at how asymmetry between competitive effect and response changes with nitrogen addition. We found that nitrogen did increase response-effect asymmetry in networks composed only of slow growing species, mostly due to increased competitive effects of species such as *Daucus carota, Prunella grandiflora* and *Plantago media* (Fig. S2), and not due to decreases in their responses. However, we observed the opposite outcome in fast growing and mixed fast-slow communities, where nitrogen led to more even and symmetrical competition networks. Nitrogen only increased competitive effects for a few fast-growing species, like *Gallium album*, while it reduced some extreme competitive effects and increased the competitive response of many others, such as *Holcus lanatus* (Fig. S2). These differences between slow and fast networks might have several explanations: adding nitrogen alone could shift resource limitation to phosphorus or water, rather than to light (Li, Niu, and Yu 2016; Lü et al. 2018; Dong et al. 2019), and slow growing species may be very competitive for phosphorus or water. In addition, several fast growing species may be very uncompetitive without nitrogen, leading to highly uneven competition networks under resource limited conditions. Although more work is needed to test these possibilities, these network changes align with the fact that adding nitrogen shifts our experimental communities toward lower values of SLA over time, which suggests that slow species are more competitive on N fertilised plots (Fig. S8). Overall, these results indicate that species with different growth strategies may compete in fundamentally different ways with consequences for how their competition networks change in response to resource addition.

Removal of foliar fungal pathogens had a large effect on our competition networks but the effects contradicted our main hypotheses (Table 1). Fungicide addition reduced both negative skewness (fewer extreme values) and modularity and increased the evenness of interaction coefficients (Fig. 3). In other words, it made competition among species more similar. The increase in evenness occurred in all networks and seemed to be caused by the fact that fungal pathogens reduced growth of weak competitors, as these species strongly increased in cover when fungicide was applied (Fig. S9). However, the decrease in skewness and modularity was more pronounced in slow only, or mixed fast-slow networks. Fast growing species are more affected by pathogens (Cappelli et al. 2020) and may have become more competitive against slow species with fungicide, which would make mixed networks less skewed and modular. In contrast to their destabilising effects, by making competition more unequal, foliar fungal pathogens did increase intraspecific competition, which should stabilise the dynamics of interacting species. This agrees with the large literature suggesting that pathogens drive stabilising Janzen-Connell effects (Adler and Muller-Landau 2005; Comita et al. 2014; Bagchi et al. 2014; Liu et al. 2022) Therefore, our results collectively indicate that pathogens can destabilise communities by targeting fast-growing species, that are weak competitors in our experiment, but they can also stabilize communities by increasing conspecific negative density dependence.

We found strong interactions between nitrogen and fungicide on multiple network metrics. We expected nitrogen and fungicide to have similar effects and to both promote dominance and reduce evenness (Mitchell et al. 2003). However, this was not the case and nitrogen and fungicide typically dampened each other’s effects (Fig. 3). We initially thought this might be because nitrogen and fungicide each favoured a different set of species, i.e., some species became more competitive with nitrogen and others with fungicide. However, it seems instead that individual pairwise competition coefficients changed in different ways in response to nitrogen versus fungicide (see highlighted examples in Fig. S3). This often led to similar competitive effects for individual species in the control and combined treatment, for example, both nitrogen and fungicide alone increased the competitive effects of the weakest competitor (*Crepis biennis*) and decreased the competitive effects of the strongest competitor (*Holcus lanatus*) but both species had similar competitive effects in the control and combined treatments (Fig. S2). Contrary to Wang et al. (2010), we found that overall, our treatments altered competitive effects more than responses. Competitive response is directly related to species persistence over time (Godoy, Kraft, and Levine 2014), so competitive responses may be selected to be more stable, while competitive effects could vary more with environmental conditions. Our results show that resource addition and enemy removal can interact in complex ways to affect competition and they highlight the value of characterising competition at a pairwise level.

Seasonality had a comparatively small impact on network metrics, indicating that our results are robust and not driven by responses measured at a single time point (Fig. 3). Modularity was higher overall in August, which hints toward different growth strategies in our experiment during drier months, perhaps because plants suffer from more resource and water limitation (Grime 2006; Fischer et al. 2015). Interestingly, we also found an interaction effect between SLA variance and seasonality, as mixed communities were more even in June, and less even in August (Fig. S5). It therefore seems that mixing functional strategies could positively impact diversity in June, but negatively in August. Field management (cutting in mid-June) could explain the negative impact in the second half of the growing season, as mowing might favour species with higher disturbance tolerance and ability to resprout (Bellingham and Sparrow 2000).

Having characterised how the structure of species interactions changes with resources and enemies, the next step would be to assess the system dynamics, i.e., which growth strategies will decline or go extinct (losers), and which will increase and become dominant (winners). We could not rigorously evaluate whether less diagonally dominant and more negatively skewed, asymmetric and uneven structures, which we observed for slow species under nitrogen addition, maintain lower diversity. Nevertheless, we have observed that the PaNDiv communities treated with nitrogen are shifting toward lower values of SLA (Fig. S8), suggesting that slow species are more competitive with nitrogen, and we also found a relationship between cover change in fungicide plots and species competitive effects (Fig. S9). Therefore, these preliminary data may indicate an empirical connection between the structure of the interactions and shifts in species abundance. We only modelled change in biomass of our phytometers in response to the cover of their neighbours. This assumes that competitive interactions do not change with ontogeny, thus, distinguishing life history stages (e.g., juveniles versus adults) is not important for understanding population dynamics. Previous work has shown the importance of life-stage dynamics for plant-plant interactions (Schiffers and Tielbörger 2006; Kinlock 2021); however, in order to gain generalisation and make it feasible to measure multiple species in different treatments, we have followed the study by Cardinaux, Hart and Alexander (2018), and used the simplest approach to modelling perennial plant population dynamics. Further work is needed to better test how this predicts long-term changes in population dynamics.

In conclusion, we show that resources and enemies have large impacts on plant-plant interaction networks, and importantly, these impacts depend on plants fast-slow growth strategy. These results suggest that incorporating trait-based approaches into the study of species interactions provide a way to systematically scale-up from individual species responses to changes at the entire community level. This was an aim recently highlighted in the literature that remains poorly addressed (Levine et al. 2017; Losapio, Montesinos-Navarro, and Saiz 2019). Our result further show that species with different growth strategies might interact and coexist in fundamentally different ways, which is an idea that has not been typically considered by studies linking traits to competition, which have tended to focus more on trait differences to predict competition (Adler et al. 2018). Quantifying all pairwise interactions in highly diverse ecosystems across contrasted environmental conditions is extremely challenging. However, our result shows that this effort is worthwhile to better mechanistically understand how nitrogen addition and pathogen removal, two of the most common drivers of global change, affect complex communities. Taken together, our work highlights the necessity of combining a network and a trait-based perspective to progress in our understanding of the effects of global change drivers on diverse plant communities.

## Acknowledgements

We are grateful to the whole PaNDiv team, especially Sylvain Chartier, Mervi Laitinen, Hugo Vincent and many helpers, for maintaining the experiment. We also thank Seraina Cappelli and Noémie Pichon for their role in setting up the PaNDiv experiment. Several people helped with collecting the plant cover data, in particular: Vera Alessandrello, Hannah Bratschi, Eli Bucher, Tala Bürki, Géraldine Chavey, Chiara Durrer, Matthieu Gauvrit, Benjamin Herren, Fabian Heussler, Vinciane Horner, Sandy Kalaydjian, Nadia Maaroufi, Olivier Magnin, Dmitry Maryasov, Anja Michel, Thu Zar Nwe, Barryette Oberholzer, Scarlett Peréz Gordillo, Valentin Pulver, Georges Saumier, Nynke Van Duijin, Joseph Volery, and Lia Zehnder. The project was funded by the Swiss National Science Foundation (Award 310030_185260). Oscar Godoy acknowledges financial support provided by the Spanish Ministry of Economy and Competitiveness (MINECO) and by the European Social Fund through the Ramón y Cajal Program (RYC2017-23666). Hugo Saíz is supported by a María Zambrano fellowship funded by the Ministry of Universities and European Union-Next Generation plan.

## Author contribution

EA designed the PaNDiv Experiment and obtained the necessary funding, OG, EA and CD developed the ideas for this study, CD collected the data in PaNDiv experiment, analysed the data and wrote the manuscript with substantial inputs from OG, EA and HS. All authors contributed substantially to revisions.

## Data accessibility

All data and R code is available in a public GitHub repository at this address: https://github.com/cardaips/PaNDiv_competition_networks, a webpage deployment is also available at this address: https://cardaips.github.io/PaNDiv_competition_networks.

## Box 1: Equations for calculating network metrics

With the matrices of species interactions obtained in the previous step, we calculated the following metrics at the community level (Table 1):

> - *The global ratio of intraspecific versus interspecific competition (D)*

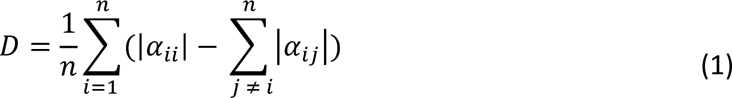

Where *D* is a measure of the diagonal dominance of the matrix, *n* is the total number of species in the matrix, α_*ii*_ is the intraspecific competition coefficient (contained in the diagonal), and α_*ij*_ is the interspecific effect of species j on speciesi (contained in the off diagonal). If *D* ≥ 0, then species suffer stronger intraspecific competition than the sum of all possible interspecific competition coefficients. Reciprocally, if *D* ≤ 0, the sum of interspecific competition is larger than intraspecific competition. This continuous measure is conceptually close to the binary measure of quasi diagonal dominance (QDD) which has been related to network stability (Liang and Wu 1998).

> - *The degree of asymmetry (A) in competition*

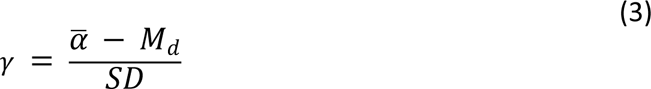

Where *A* measures the difference between the competitive effect of species i on species j (α_*ij*_), versus the effect of j on i (α_*ji*_), averaged across all pairs of species (Vázquez et al. 2007). A greater value of *A* implies larger asymmetry within the community.

> - *The distribution of interaction coefficients measured as skewness (*γ)

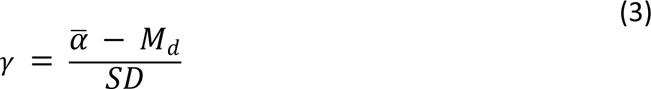

Where 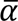 is the mean of all interaction coefficients α_*ij*_ within the network, *M*_*d*_ the median, and *SD* the standard deviation of the coefficients. The skewness, by comparing the mean to the median, describes to the extent to which interaction coefficients are skewed to either negative or positive values, i.e., if there are some extremely low or high values. Greater skewness values indicate a distribution of interaction coefficients with larger tails.

> - The distribution of interaction coefficients measured as the Gini coefficient of evenness (G),

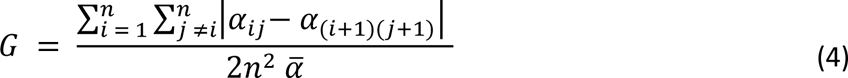

Where *n* is the number of species, <u>α</u> is the mean of every interaction coefficient α_*ij*_. The Gini coefficient describes the degree of inequality in the distribution of the interaction coefficients. It ranges between 0 and 1 where 0 is perfect equality and 1 is total inequality (Gini 1912), we use 1-G here so that high values indicate high evenness. Because interspecific interactions coefficients were both positive and negative, we used a correction of G which separates positive and negative values, then reassembles them in a single value of evenness (Raffinetti, Siletti, and Vernizzi 2015).

> - *The modularity (M)*

Modularity (M) is computed as a ratio that compares the average strength of the links between two nodes, or vertex, that belong to the same module, versus links between two nodes of different modules. Consequently, optimising modularity corresponds to grouping the nodes, i.e. the species, to maximise the strength of interactions (interaction coefficients are used as weights) within a module, while minimising the strength of interactions between modules. This creates the smallest number of groups of similarly behaving species. Modularity was computed using absolute values of interaction coefficients, following the formula from Clauset, Newman, and Moore 2004:

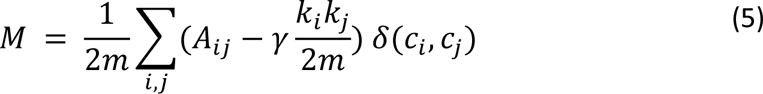

Where *m* is the number of edges, *A*_*ij*_ is the adjacency matrix (weighted by the magnitude of their corresponding interactions α_*ij*_), γ is the resolution parameter weighing for the size of the clusters (γ = 1 here, as standard), *k*_*i*_ is the degree of connection going out from the node, *k*_j_ the one coming to the node, δ(*c*_*i*_, *c_j_*) is equal to 1 if both vertex are belonging to the same cluster, 0 otherwise. By weighing the adjacency matrix by the interaction magnitude and not sign, we assume that a strong positive link is of the same importance as a strong negative link. Negative values of modularity mean that interactions are stronger between than within modules, while positive values indicate the opposite (Clauset, Newman, and Moore 2004; Reichardt and Bornholdt 2006).

## Notes

### Competing Interest Statement

The authors have declared no competing interest.

## References

Adler, Frederick R., and Helene C. Muller-Landau. 2005. ‘When Do Localized Natural Enemies Increase Species Richness?’ Ecology Letters 8 (4): 438–47. 10.1111/j.1461-0248.2005.00741.x.

Adler, Peter B., Danielle Smull, Karen H. Beard, Ryan T. Choi, Tucker Furniss, Andrew Kulmatiski, Joan M. Meiners, Andrew T. Tredennick, and Kari E. Veblen. 2018. ‘Competition and Coexistence in Plant Communities: Intraspecific Competition Is Stronger than Interspecific Competition’. Ecology Letters 21 (9): 1319–29. 10.1111/ele.13098.

Ahmad, Altaf, Hema Diwan, and Yash P. Abrol. 2010. ‘Global Climate Change, Stress and Plant Productivity’. In Abiotic Stress Adaptation in Plants: Physiological, Molecular and Genomic Foundation, edited by Ashwani Pareek, S.K. Sopory, and Hans J. Bohnert, 503–21. Dordrecht: Springer Netherlands. 10.1007/978-90-481-3112-9_23.

Alexander, Helen M., and Robert D. Holt. 1998. ‘The Interaction between Plant Competition and Disease’. Perspectives in Plant Ecology, Evolution and Systematics 1 (2): 206–20. 10.1078/1433-8319-00059.

Allan, Eric, and Michael J. Crawley. 2011. ‘Contrasting Effects of Insect and Molluscan Herbivores on Plant Diversity in a Long-Term Field Experiment’. Ecology Letters 14 (12): 1246–53. 10.1111/j.1461-0248.2011.01694.x.

Allan, Eric, Jasper van Ruijven, and Michael J. Crawley. 2010. ‘Foliar Fungal Pathogens and Grassland Biodiversity’. Ecology 91 (9): 2572–82. 10.1890/09-0859.1.

Allesina, Stefano, and Jonathan M. Levine. 2011. ‘A Competitive Network Theory of Species Diversity’. Proceedings of the National Academy of Sciences 108 (14): 5638–42. 10.1073/pnas.1014428108.

Bagchi, Robert, Rachel E. Gallery, Sofia Gripenberg, Sarah J. Gurr, Lakshmi Narayan, Claire E. Addis, Robert P. Freckleton, and Owen T. Lewis. 2014. ‘Pathogens and Insect Herbivores Drive Rainforest Plant Diversity and Composition’. Nature 506 (7486): 85–88. 10.1038/nature12911.

Bates, Douglas, Martin Mächler, Ben Bolker, and Steve Walker. 2015. ‘Fitting Linear Mixed-Effects Models Using Lme4’. Journal of Statistical Software 67 (October): 1–48. 10.18637/jss.v067.i01.

Bellingham, Peter J., and Ashley D. Sparrow. 2000. ‘Resprouting as a Life History Strategy in Woody Plant Communities’. Oikos 89 (2): 409–16. 10.1034/j.1600-0706.2000.890224.x.

Bimler, Malyon D., Daniel B. Stouffer, Hao Ran Lai, and Margaret M. Mayfield. 2018. ‘Accurate Predictions of Coexistence in Natural Systems Require the Inclusion of Facilitative Interactions and Environmental Dependency’. Journal of Ecology 106 (5): 1839–52. 10.1111/1365-2745.13030.

Blüthgen, Nico, Carsten F. Dormann, Daniel Prati, Valentin H. Klaus, Till Kleinebecker, Norbert Hölzel, Fabian Alt, et al. 2012. ‘A Quantitative Index of Land-Use Intensity in Grasslands: Integrating Mowing, Grazing and Fertilization’. Basic and Applied Ecology 13 (3): 207–20. 10.1016/j.baae.2012.04.001.

Brooker, Rob W. 2006. ‘Plant–Plant Interactions and Environmental Change’. New Phytologist 171 (2): 271–84. 10.1111/j.1469-8137.2006.01752.x.

Brooks, Mollie E., Kasper Kristensen, Koen J. van Benthem, Arni Magnusson, Casper W. Berg, Anders Nielsen, Hans J. Skaug, Martin Mächler, and Benjamin M. Bolker. 2017. ‘glmmTMB Balances Speed and Flexibility Among Packages for Zero-Inflated Generalized Linear Mixed Modeling’. The R Journal 9 (2): 378–400.

Buche, Lisa, Jurg W. Spaak, Javier Jarillo, and Frederik De Laender. 2022. ‘Niche Differences, Not Fitness Differences, Explain Predicted Coexistence across Ecological Groups’. Journal of Ecology 110 (11): 2785–96. 10.1111/1365-2745.13992.

Cappelli, Seraina L., Noémie A. Pichon, Anne Kempel, and Eric Allan. 2020. ‘Sick Plants in Grassland Communities: A Growth-Defense Trade-off Is the Main Driver of Fungal Pathogen Abundance’. Ecology Letters 23 (9): 1349–59. 10.1111/ele.13537.

Cappelli, Seraina L., Noémie A. Pichon, Tosca Mannall, and Eric Allan. 2022. ‘Partitioning the Effects of Plant Diversity on Ecosystem Functions at Different Trophic Levels’. Ecological Monographs 92 (3): e1521. 10.1002/ecm.1521.

Cardinaux, Aline, Simon P. Hart, and Jake M. Alexander. 2018. ‘Do Soil Biota Influence the Outcome of Novel Interactions between Plant Competitors?’ Journal of Ecology 106 (5): 1853–63. 10.1111/1365-2745.13029.

Chesson, Peter. 2000. ‘Mechanisms of Maintenance of Species Diversity’. Annual Review of Ecology and Systematics 31 (1): 343–66.

Chesson, Peter, and Jessica J. Kuang. 2008. ‘The Interaction between Predation and Competition’. Nature 456 (7219): 235–38. 10.1038/nature07248.

Clauset, Aaron, M. E. J. Newman, and Cristopher Moore. 2004. ‘Finding Community Structure in Very Large Networks’. Physical Review E 70 (6): 066111. 10.1103/PhysRevE.70.066111.

Cleland, Elsa E., and W. Stanley Harpole. 2010. ‘Nitrogen Enrichment and Plant Communities’. Annals of the New York Academy of Sciences 1195 (1): 46–61. 10.1111/j.1749-6632.2010.05458.x.

Coley, Phyllis D., John P. Bryant, and F. Stuart Chapin. 1985. ‘Resource Availability and Plant Antiherbivore Defense’. Science 230 (4728): 895–99. 10.1126/science.230.4728.895.

Comita, Liza S., Simon A. Queenborough, Stephen J. Murphy, Jenalle L. Eck, Kaiyang Xu, Meghna Krishnadas, Noelle Beckman, and Yan Zhu. 2014. ‘Testing Predictions of the Janzen–Connell Hypothesis: A Meta-Analysis of Experimental Evidence for Distance- and Density-Dependent Seed and Seedling Survival’. Journal of Ecology 102 (4): 845–56. 10.1111/1365-2745.12232.

Connell, Joseph H. 1971. ‘On the Role of Natural Enemies in Preventing Competitive Exclusion in Some Marine Animals and in Rain Forest Trees’. Dynamics of Populations 298: 312.

Crawley, M. J., A. E. Johnston, J. Silvertown, M. Dodd, C. de Mazancourt, M. S. Heard, D. F. Henman, and G. R. Edwards. 2005. ‘Determinants of Species Richness in the Park Grass Experiment.’ The American Naturalist 165 (2): 179–92. 10.1086/427270.

Csardi, Maintainer Gabor. 2013. ‘Package “Igraph”’. Last Accessed 3 (09): 2013.

DeMalach, Niv, Eli Zaady, and Ronen Kadmon. 2017. ‘Light Asymmetry Explains the Effect of Nutrient Enrichment on Grassland Diversity’. Ecology Letters 20 (1): 60–69.

Díaz, Sandra, Jens Kattge, Johannes H. C. Cornelissen, Ian J. Wright, Sandra Lavorel, Stéphane Dray, Björn Reu, et al. 2016. ‘The Global Spectrum of Plant Form and Function’. Nature 529 (7585): 167–71. 10.1038/nature16489.

DiTommaso, Antonio, and L. W. Aarssen. 1989. ‘Resource Manipulations in Natural Vegetation: A Review’. Vegetatio 84 (1): 9–29. 10.1007/BF00054662.

Dong, Chengcheng, Wei Wang, Hongyan Liu, Xiaotian Xu, and Hui Zeng. 2019. ‘Temperate Grassland Shifted from Nitrogen to Phosphorus Limitation Induced by Degradation and Nitrogen Deposition: Evidence from Soil Extracellular Enzyme Stoichiometry’. Ecological Indicators 101 (June): 453–64. 10.1016/j.ecolind.2019.01.046.

Dormann, Carsten F., and Stephen H. Roxburgh. 2005. ‘Experimental Evidence Rejects Pairwise Modelling Approach to Coexistence in Plant Communities’. Proceedings of the Royal Society B: Biological Sciences 272 (1569): 1279–85.

Eskelinen, Anu, W. Stanley Harpole, Maria-Theresa Jessen, Risto Virtanen, and Yann Hautier. 2022. ‘Light Competition Drives Herbivore and Nutrient Effects on Plant Diversity’. Nature 611 (7935): 301–5. 10.1038/s41586-022-05383-9.

Fischer, A. M., D. E. Keller, M. A. Liniger, J. Rajczak, C. Schär, and C. Appenzeller. 2015. ‘Projected Changes in Precipitation Intensity and Frequency in Switzerland: A Multi-Model Perspective’. International Journal of Climatology 35 (11): 3204–19. 10.1002/joc.4162.

Funk, Jennifer L., Julie E. Larson, Gregory M. Ames, Bradley J. Butterfield, Jeannine Cavender-Bares, Jennifer Firn, Daniel C. Laughlin, Ariana E. Sutton-Grier, Laura Williams, and Justin Wright. 2017. ‘Revisiting the Holy Grail: Using Plant Functional Traits to Understand Ecological Processes’. Biological Reviews 92 (2): 1156–73. 10.1111/brv.12275.

Gallien, Laure, Niklaus E. Zimmermann, Jonathan M. Levine, and Peter B. Adler. 2017. ‘The Effects of Intransitive Competition on Coexistence’. Ecology Letters 20 (7): 791–800. 10.1111/ele.12775.

Gini, C. 1912. ‘Variabilitá e Mutabilitá, Con-Tributo Allo Studio Delle Distribuzioni: Relazioni Statistische’. Studi Economico-Guiridici Della R. Universitá Di Cagliari.

Godoy, Oscar, Nathan J. B. Kraft, and Jonathan M. Levine. 2014. ‘Phylogenetic Relatedness and the Determinants of Competitive Outcomes’. Ecology Letters 17 (7): 836–44. 10.1111/ele.12289.

Goldberg, Deborah E., and Keith Landa. 1991. ‘Competitive Effect and Response: Hierarchies and Correlated Traits in the Early Stages of Competition’. Journal of Ecology 79 (4): 1013–30. 10.2307/2261095.

Granjel, Rodrigo R., Eric Allan, and Oscar Godoy. 2023. ‘Nitrogen Enrichment and Foliar Fungal Pathogens Affect the Mechanisms of Multispecies Plant Coexistence’. New Phytologist 237 (6): 2332–46.

Grigulis, Karl, Sandra Lavorel, Ute Krainer, Nicolas Legay, Catherine Baxendale, Maxime Dumont, Eva Kastl, et al. 2013. ‘Relative Contributions of Plant Traits and Soil Microbial Properties to Mountain Grassland Ecosystem Services’. Journal of Ecology 101 (1): 47–57. 10.1111/1365-2745.12014.

Grime, J. Philip. 2006. Plant Strategies, Vegetation Processes, and Ecosystem Properties. John Wiley & Sons.

Grime, J.P. 1979. ‘Primary Strategies in Plants’. Transactions of the Botanical Society of Edinburgh 43 (2): 151–60. 10.1080/03746607908685348.

Gross, Nicolas, Georges Kunstler, Pierre Liancourt, Francesco De Bello, Katharine Nash Suding, and Sandra Lavorel. 2009. ‘Linking Individual Response to Biotic Interactions with Community Structure: A Trait-Based Framework’. Functional Ecology 23 (6): 1167–78. 10.1111/j.1365-2435.2009.01591.x.

Hadfield, Jarrod D. 2010. ‘MCMC Methods for Multi-Response Generalized Linear Mixed Models: The MCMCglmm R Package’. Journal of Statistical Software 33 (February): 1–22. 10.18637/jss.v033.i02.

Harpole, W. Stanley, Lauren L. Sullivan, Eric M. Lind, Jennifer Firn, Peter B. Adler, Elizabeth T. Borer, Jonathan Chase, et al. 2016. ‘Addition of Multiple Limiting Resources Reduces Grassland Diversity’. Nature 537 (7618): 93–96. 10.1038/nature19324.

Hautier, Yann, Pascal A. Niklaus, and Andy Hector. 2009. ‘Competition for Light Causes Plant Biodiversity Loss After Eutrophication’. Science 324 (5927): 636–38. 10.1126/science.1169640.

Holt, Robert D. 1977. ‘Predation, Apparent Competition, and the Structure of Prey Communities’. Theoretical Population Biology 12 (2): 197–229. 10.1016/0040-5809(77)90042-9.

Holt, Robert D., James Grover, and David Tilman. 1994. ‘Simple Rules for Interspecific Dominance in Systems with Exploitative and Apparent Competition’. The American Naturalist 144 (5): 741–71. 10.1086/285705.

Isbell, Forest, Peter B. Reich, David Tilman, Sarah E. Hobbie, Stephen Polasky, and Seth Binder. 2013. ‘Nutrient Enrichment, Biodiversity Loss, and Consequent Declines in Ecosystem Productivity’. Proceedings of the National Academy of Sciences 110 (29): 11911–16.

Janzen, Daniel H. 1970. ‘Herbivores and the Number of Tree Species in Tropical Forests’. The American Naturalist 104 (940): 501–28. 10.1086/282687.

Kinlock, Nicole L. 2019. ‘A Meta-Analysis of Plant Interaction Networks Reveals Competitive Hierarchies as Well as Facilitation and Intransitivity’. The American Naturalist 194 (5): 640–53. 10.1086/705293.

Kinlock, Nicole L. 2021. ‘Uncovering Structural Features That Underlie Coexistence in an Invaded Woody Plant Community with Interaction Networks at Multiple Life Stages’. Journal of Ecology 109 (1): 384–98. 10.1111/1365-2745.13489.

Komsta, Lukasz, and Frederick Novomestky. 2015. ‘Moments, Cumulants, Skewness, Kurtosis and Related Tests’. R Package Version 14.

Kraft, Nathan J. B., Oscar Godoy, and Jonathan M. Levine. 2015. ‘Plant Functional Traits and the Multidimensional Nature of Species Coexistence’. Proceedings of the National Academy of Sciences 112 (3): 797–802. 10.1073/pnas.1413650112.

Kunstler, Georges, Daniel Falster, David A. Coomes, Francis Hui, Robert M. Kooyman, Daniel C. Laughlin, Lourens Poorter, et al. 2016. ‘Plant Functional Traits Have Globally Consistent Effects on Competition’. Nature 529 (7585): 204–7. 10.1038/nature16476.

Levine, Jonathan M., Jordi Bascompte, Peter B. Adler, and Stefano Allesina. 2017. ‘Beyond Pairwise Mechanisms of Species Coexistence in Complex Communities’. Nature 546 (7656): 56–64. 10.1038/nature22898.

Li, Yong, Shuli Niu, and Guirui Yu. 2016. ‘Aggravated Phosphorus Limitation on Biomass Production under Increasing Nitrogen Loading: A Meta-Analysis’. Global Change Biology 22 (2): 934–43. 10.1111/gcb.13125.

Liang, Xue-Bin, and Li-De Wu. 1998. ‘New Sufficient Conditions for Absolute Stability of Neural Networks’. IEEE Transactions on Circuits and Systems I: Fundamental Theory and Applications 45 (5): 584–86. 10.1109/81.668873.

Liu, Xiang, Ingrid M. Parker, Gregory S. Gilbert, Yawen Lu, Yao Xiao, Li Zhang, Mengjiao Huang, Yikang Cheng, Zhenhua Zhang, and Shurong Zhou. 2022. ‘Coexistence Is Stabilized by Conspecific Negative Density Dependence via Fungal Pathogens More than Oomycete Pathogens’. Ecology 103 (12): e3841. 10.1002/ecy.3841.

López-Angulo, Jesús, Nathan G. Swenson, Lohengrin A. Cavieres, and Adrián Escudero. 2018. ‘Interactions between Abiotic Gradients Determine Functional and Phylogenetic Diversity Patterns in Mediterranean-Type Climate Mountains in the Andes’. Journal of Vegetation Science 29 (2): 245–54. 10.1111/jvs.12607.

Loranger, Jessy, Sebastian T. Meyer, Bill Shipley, Jens Kattge, Hannah Loranger, Christiane Roscher, and Wolfgang W. Weisser. 2012. ‘Predicting Invertebrate Herbivory from Plant Traits: Evidence from 51 Grassland Species in Experimental Monocultures’. Ecology 93 (12): 2674–82. 10.1890/12-0328.1.

Loreau, Michel, and Andy Hector. 2001. ‘Partitioning Selection and Complementarity in Biodiversity Experiments’. Nature 412 (6842): 72–76. 10.1038/35083573.

Losapio, Gianalberto, Alicia Montesinos-Navarro, and Hugo Saiz. 2019. ‘Perspectives for Ecological Networks in Plant Ecology’. Plant Ecology & Diversity 12 (2): 87–102. 10.1080/17550874.2019.1626509.

Losapio, Gianalberto, Christian Schöb, Phillip P. A. Staniczenko, Francesco Carrara, Gian Marco Palamara, Consuelo M. De Moraes, Mark C. Mescher, et al. 2021. ‘Network Motifs Involving Both Competition and Facilitation Predict Biodiversity in Alpine Plant Communities’. Proceedings of the National Academy of Sciences 118 (6): e2005759118. 10.1073/pnas.2005759118.

Lü, Xiao-Tao, Zhuo-Yi Liu, Yan-Yu Hu, and Hai-Yang Zhang. 2018. ‘Testing Nitrogen and Water Co-Limitation of Primary Productivity in a Temperate Steppe’. Plant and Soil 432 (1): 119–27. 10.1007/s11104-018-3791-6.

Matías, Luis, Oscar Godoy, Lorena Gómez-Aparicio, and Ignacio M. Pérez-Ramos. 2018. ‘An Experimental Extreme Drought Reduces the Likelihood of Species to Coexist despite Increasing Intransitivity in Competitive Networks’. Journal of Ecology 106 (3): 826–37. 10.1111/1365-2745.12962.

Mayfield, Margaret M., and Jonathan M. Levine. 2010. ‘Opposing Effects of Competitive Exclusion on the Phylogenetic Structure of Communities’. Ecology Letters 13 (9): 1085–93. 10.1111/j.1461-0248.2010.01509.x.

Mitchell, Charles E., Peter B. Reich, David Tilman, and James V. Groth. 2003. ‘Effects of Elevated CO2, Nitrogen Deposition, and Decreased Species Diversity on Foliar Fungal Plant Disease’. Global Change Biology 9 (3): 438–51. 10.1046/j.1365-2486.2003.00602.x.

Mordecai, Erin A. 2011. ‘Pathogen Impacts on Plant Communities: Unifying Theory, Concepts, and Empirical Work’. Ecological Monographs 81 (3): 429–41. 10.1890/10-2241.1.

Pacala, S. W., and M. J. Crawley. 1992. ‘Herbivores and Plant Diversity’. The American Naturalist 140 (2): 243–60. 10.1086/285411.

Parker, Ingrid M., and Gregory S. Gilbert. 2018. ‘Density-Dependent Disease, Life-History Trade-Offs, and the Effect of Leaf Pathogens on a Suite of Co-Occurring Close Relatives’. Journal of Ecology 106 (5): 1829–38. 10.1111/1365-2745.13024.

Pichon, Noémie A., Seraina L. Cappelli, Santiago Soliveres, Norbert Hölzel, Valentin H. Klaus, Till Kleinebecker, and Eric Allan. 2020. ‘Decomposition Disentangled: A Test of the Multiple Mechanisms by Which Nitrogen Enrichment Alters Litter Decomposition’. Functional Ecology 34 (7): 1485–96. 10.1111/1365-2435.13560.

Poorter, Hendrik, Carlo Remkes, and Hans Lambers. 1990. ‘Carbon and Nitrogen Economy of 24 Wild Species Differing in Relative Growth Rate’. Plant Physiology 94 (2): 621–27. 10.1104/pp.94.2.621.

Raffinetti, Emanuela, Elena Siletti, and Achille Vernizzi. 2015. ‘On the Gini Coefficient Normalization When Attributes with Negative Values Are Considered’. Statistical Methods & Applications 24 (3): 507–21. 10.1007/s10260-014-0293-4.

Reichardt, Jörg, and Stefan Bornholdt. 2006. ‘Statistical Mechanics of Community Detection’. Physical Review E 74 (1): 016110. 10.1103/PhysRevE.74.016110.

Saiz, Hugo, Yoann Le Bagousse-Pinguet, Nicolas Gross, and Fernando T. Maestre. 2019. ‘Intransitivity Increases Plant Functional Diversity by Limiting Dominance in Drylands Worldwide’. Journal of Ecology 107 (1): 240–52. 10.1111/1365-2745.13018.

Schiffers, Katja, and Katja Tielbörger. 2006. ‘Ontogenetic Shifts in Interactions among Annual Plants’. Journal of Ecology 94 (2): 336–41.

Schwinning, Susanne, and Jacob Weiner. 1998. ‘Mechanisms Determining the Degree of Size Asymmetry in Competition among Plants’. Oecologia 113 (4): 447–55. 10.1007/s004420050397.

Silveira Pontes, Laíse da, Frédérique Louault, Pascal Carrère, Vincent Maire, Donato Andueza, and Jean-François Soussana. 2010. ‘The Role of Plant Traits and Their Plasticity in the Response of Pasture Grasses to Nutrients and Cutting Frequency’. Annals of Botany 105 (6): 957–65. 10.1093/aob/mcq066.

Soliveres, Santiago, Fernando T. Maestre, Werner Ulrich, Peter Manning, Steffen Boch, Matthew A. Bowker, Daniel Prati, et al. 2015. ‘Intransitive Competition Is Widespread in Plant Communities and Maintains Their Species Richness’. Ecology Letters 18 (8): 790–98. 10.1111/ele.12456.

Spear, Erin R., and Erin A. Mordecai. 2018. ‘Foliar Pathogens Are Unlikely to Stabilize Coexistence of Competing Species in a California Grassland’. Ecology 99 (10): 2250–59. 10.1002/ecy.2427.

Suding, Katharine N., Scott L. Collins, Laura Gough, Christopher Clark, Elsa E. Cleland, Katherine L. Gross, Daniel G. Milchunas, and Steven Pennings. 2005. Proceedings of the National Academy of Sciences 102 (12): 4387–92.

Tilman, David. 1982. Resource Competition and Community Structure. (MPB-17), Volume 17. Resource Competition and Community Structure. (MPB-17), *Volume* 17. Princeton University Press. 10.1515/9780691209654.

Tilman, David. 1985. ‘The Resource-Ratio Hypothesis of Plant Succession’. The American Naturalist 125 (6): 827–52. 10.1086/284382.

Van Dyke, Mary N., Jonathan M. Levine, and Nathan J. B. Kraft. 2022. ‘Small Rainfall Changes Drive Substantial Changes in Plant Coexistence’. Nature 611 (7936): 507–11. 10.1038/s41586-022-05391-9.

Vázquez, Diego P., Carlos J. Melián, Neal M. Williams, Nico Blüthgen, Boris R. Krasnov, and Robert Poulin. 2007. ‘Species Abundance and Asymmetric Interaction Strength in Ecological Networks’. Oikos 116 (7): 1120–27. 10.1111/j.0030-1299.2007.15828.x.

Wang, Ping, Tara Stieglitz, Dao Wei Zhou, and James F Cahill Jr. 2010. ‘Are Competitive Effect and Response Two Sides of the Same Coin, or Fundamentally Different?’ Functional Ecology 24 (1): 196–207. 10.1111/j.1365-2435.2009.01612.x.

Wickham, Hadley, Jim Hester, Winston Chang, and Maintainer Jim Hester. 2021. Package ‘Devtools’.

Xiao, Yao, Xiang Liu, Li Zhang, Zhiping Song, and Shurong Zhou. 2021. ‘The Allometry of Plant Height Explains Species Loss under Nitrogen Addition’. Ecology Letters 24 (3): 553–62. 10.1111/ele.13673.

Yang, Xuejun, Lorena Gómez-Aparicio, Christopher J. Lortie, Miguel Verdú, Lohengrin A. Cavieres, Zhenying Huang, Ruiru Gao, Rong Liu, Yonglan Zhao, and Johannes H. C. Cornelissen. 2022. ‘Net Plant Interactions Are Highly Variable and Weakly Dependent on Climate at the Global Scale’. Ecology Letters 25 (6): 1580–93. 10.1111/ele.14010.

