## Supplementary material for "Fast-slow traits predict competition network structure and its response to resources and enemies"

- Supplementary Method 1. SLA calculations

One objective of our analyses was to compare whether the structure of plant interaction networks varies between fast and slow growing species. In order to differentiate these strategies, we used values of SLA measured in June and August 2019 in PaNDiv monocultures. For each species, treatment, and sampling period, 5 leaves were sampled and their area (cm2), fresh weight (mg) and dry weight (mg) were measured. SLA (m2 kg−1) was averaged per species between all treatments and across the two sampling periods (in order to prevent any correlation between SLA and treatments). Then, mean and variance of SLA were calculated for each five species network. SLA was log transformed to reduce the correlation between the mean SLA and the variance (Pearson's correlation coefficient, c ~ 0.28 after log transformation).

- Table S1. PaNDiv experimental species categorised into functional group and growth form. *Hs* and *As* (in grey) were not included as phytometer in the experiment due to germination issues. Species marked with a * were not included in the targeted neighbourhoods, but coefficient were still derived for every pairwise interactions.

| **Plant species (abbreviation)** | **Functional group** | **Growth form** |
|---|---|---|
| *Dactylis glomerata (Dg)*<br>*Holcus lanatus (Hl)*<br>*Lolium perenne (Lp)*<br>*Poa trivialis (Pt)** | Fast growing | Grasses |
| *Anthoxanthum odoratum (Ao)**<br>*Bromus erectus (Be)*<br>*Festuca rubra (Fr)*<br>*Helictotrichon pubescens (Hp)* | Slow growing | |
| *Anthriscus sylvestris (As), not included*<br>*Crepis biennis (Cb)*<br>*Galium album (Ga)*<br>*Heracleum sphondylium (Hs), not included*<br>*Rumex acetosa (Ra)**<br>*Taraxacum officinale (To)* | Fast growing | Herbs |
| *Achillea millefolium (Am)*<br>*Centaurea jacea (Cj)*<br>*Daucus carota (Dc)*<br>*Plantago media (Pm)*<br>*Prunella grandiflora (Pg)*<br>*Salvia pratensis (Sp)* | Slow growing | |

- Table S2. Outputs from multi-membership LMER models showing the effects of functional groups and treatments on networks of 5 species richness across June and August. Each column contains the results for one model and a specific network metric (scaled between 0 and 1).

| | ***Diagonal dominance*** | ***Asymmetry*** | ***Skewness*** | ***Evenness*** | ***Modularity*** |
|---|---|---|---|---|---|
| *Nitrogen* | −0.25*** (0.000) | 0.19*** (0.000) | 0.23*** (0.000) | 0.31*** (0.000) | 0.68*** (0.000) |
| *Fungicide* | −0.28*** (0.000) | 0.00 (0.938) | 0.65*** (0.000) | 0.96*** (0.000) | −0.35*** (0.000) |
| *SLA* | 0.05 (0.792) | 0.21 (0.072) | 0.09 (0.584) | −0.17 (0.235) | −0.23* (0.021) |
| *SLA variance* | 0.03*** (0.000) | −0.05*** (0.000) | −0.16*** (0.000) | 0.10*** (0.000) | 0.34*** (0.000) |
| *Season* | −0.01** (0.008) | −0.01* (0.049) | −0.04*** (0.000) | 0.07*** (0.000) | 0.78*** (0.000) |
| *Nitrogen:Fungicide* | 0.24*** (0.000) | −0.15*** (0.000) | −0.45*** (0.000) | −0.82*** (0.000) | −0.35*** (0.000) |
| *Nitrogen:SLA* | −0.02* (0.018) | −0.25*** (0.000) | −0.26*** (0.000) | 0.09*** (0.000) | 0.04*** (0.000) |
| *Nitrogen:SLA variance* | −0.16*** (0.000) | −0.11*** (0.000) | 0.33*** (0.000) | 0.11*** (0.000) | −0.32*** (0.000) |
| *Fungicide:SLA* | 0.06*** (0.000) | −0.06*** (0.000) | −0.20*** (0.000) | 0.00 (0.981) | 0.29*** (0.000) |
| *Fungicide:SLA variance* | −0.12*** (0.000) | 0.03** (0.001) | 0.43*** (0.000) | 0.11*** (0.000) | −0.37*** (0.000) |
| *SLA:Season* | 0.10*** (0.000) | −0.09*** (0.000) | 0.08*** (0.000) | −0.06*** (0.000) | 0.10*** (0.000) |
| *SLA variance:Season* | 0.14*** (0.000) | 0.03*** (0.000) | −0.11*** (0.000) | −0.29*** (0.000) | −0.05*** (0.000) |
| *Nitrogen:Fungicide:SLA* | 0.06*** (0.000) | 0.29*** (0.000) | 0.13*** (0.000) | 0.08*** (0.000) | 0.02 (0.175) |
| *Nitrogen:Fungicide:SLA variance* | −0.04*** (0.000) | 0.15*** (0.000) | −0.36*** (0.000) | −0.09*** (0.000) | 0.17*** (0.000) |
| *SD (Observations)* | 0.62 | 0.87 | 0.70 | 0.70 | 0.71 |
| *Num.Obs.* | 68976 | 68976 | 68976 | 68976 | 68976 |
| *R2 Marg.* | 0.089 | 0.038 | 0.143 | 0.225 | 0.381 |
| *R2 Cond.* | 0.360 | 0.112 | 0.316 | 0.355 | 0.434 |

- Table S3. Outputs from multi-membership LMER models showing the effects of species richness and treatments on functionally mixed communities (5, 7, 11 and 15 species) in June. Each column contains the coefficients and corresponding p-values (in brackets) for one model and a specific network metric (scaled between 0 and 1).

| | ***Diagonal dominance*** | ***Asymmetry*** | ***Skewness*** | ***Evenness*** | ***Modularity*** |
|---|---|---|---|---|---|
| *Nitrogen* | −1.13*** (0.000) | 0.09*** (0.000) | 0.36*** (0.000) | 0.03*** (0.000) | 0.02*** (0.000) |
| *Fungicide* | −1.04*** (0.000) | 0.00 (0.999) | 0.56*** (0.000) | 0.09*** (0.000) | −0.02*** (0.000) |
| *Network size (linear)* | −12.57*** (0.000) | 0.00 (1.000) | −0.21 (0.514) | −0.02 (0.605) | −0.02* (0.024) |
| *SLA* | −0.08*** (0.000) | 0.05*** (0.000) | 0.07*** (0.000) | −0.02*** (0.000) | 0.00*** (0.000) |
| *SLA variance* | 0.27*** (0.000) | −0.01*** (0.000) | −0.12*** (0.000) | 0.00*** (0.000) | 0.01*** (0.000) |
| *Nitrogen:Fungicide* | 0.78*** (0.000) | −0.07*** (0.000) | −0.59*** (0.000) | −0.08*** (0.000) | −0.01*** (0.000) |
| *Nitrogen:Network size (linear)* | −1.18*** (0.000) | 0.00 (1.000) | 0.31*** (0.000) | 0.00 (0.230) | −0.02*** (0.000) |
| *Fungicide:Network size (linear)* | −0.94*** (0.000) | 0.00 (1.000) | 0.15*** (0.000) | 0.00 (0.870) | −0.01*** (0.000) |
| *Nitrogen:SLA* | −0.04*** (0.000) | −0.07*** (0.000) | −0.08*** (0.000) | 0.01*** (0.000) | 0.00*** (0.001) |
| *Nitrogen:SLA variance* | −0.25*** (0.000) | −0.04*** (0.000) | 0.14*** (0.000) | 0.01*** (0.000) | −0.01*** (0.000) |
| *Fungicide:SLA* | 0.10*** (0.000) | −0.02*** (0.000) | −0.04*** (0.000) | 0.00 (0.060) | 0.01*** (0.000) |
| *Fungicide:SLA variance* | −0.21*** (0.000) | 0.01*** (0.000) | 0.19*** (0.000) | 0.01*** (0.000) | −0.01*** (0.000) |
| *Nitrogen:Fungicide:Network size (linear)* | 0.63*** (0.000) | 0.00 (1.000) | −0.39*** (0.000) | −0.01*** (0.000) | 0.01*** (0.000) |
| *Nitrogen:Fungicide:SLA* | 0.13*** (0.000) | 0.08*** (0.000) | −0.02*** (0.000) | 0.01*** (0.000) | 0.00*** (0.000) |
| *Nitrogen:Fungicide:SLA variance* | −0.14*** (0.000) | 0.05*** (0.000) | −0.15*** (0.000) | −0.01*** (0.000) | 0.00*** (0.000) |
| *SD (Observations)* | 1.13 | 0.23 | 0.36 | 0.04 | 0.02 |
| *Num.Obs.* | 584’256 | 584’256 | 584’256 | 584’256 | 584’256 |
| *R2 Marg.* | 0.905 | 0.053 | 0.238 | 0.340 | 0.391 |
| *R2 Cond.* | 0.936 | 0.123 | 0.399 | 0.437 | 0.417 |

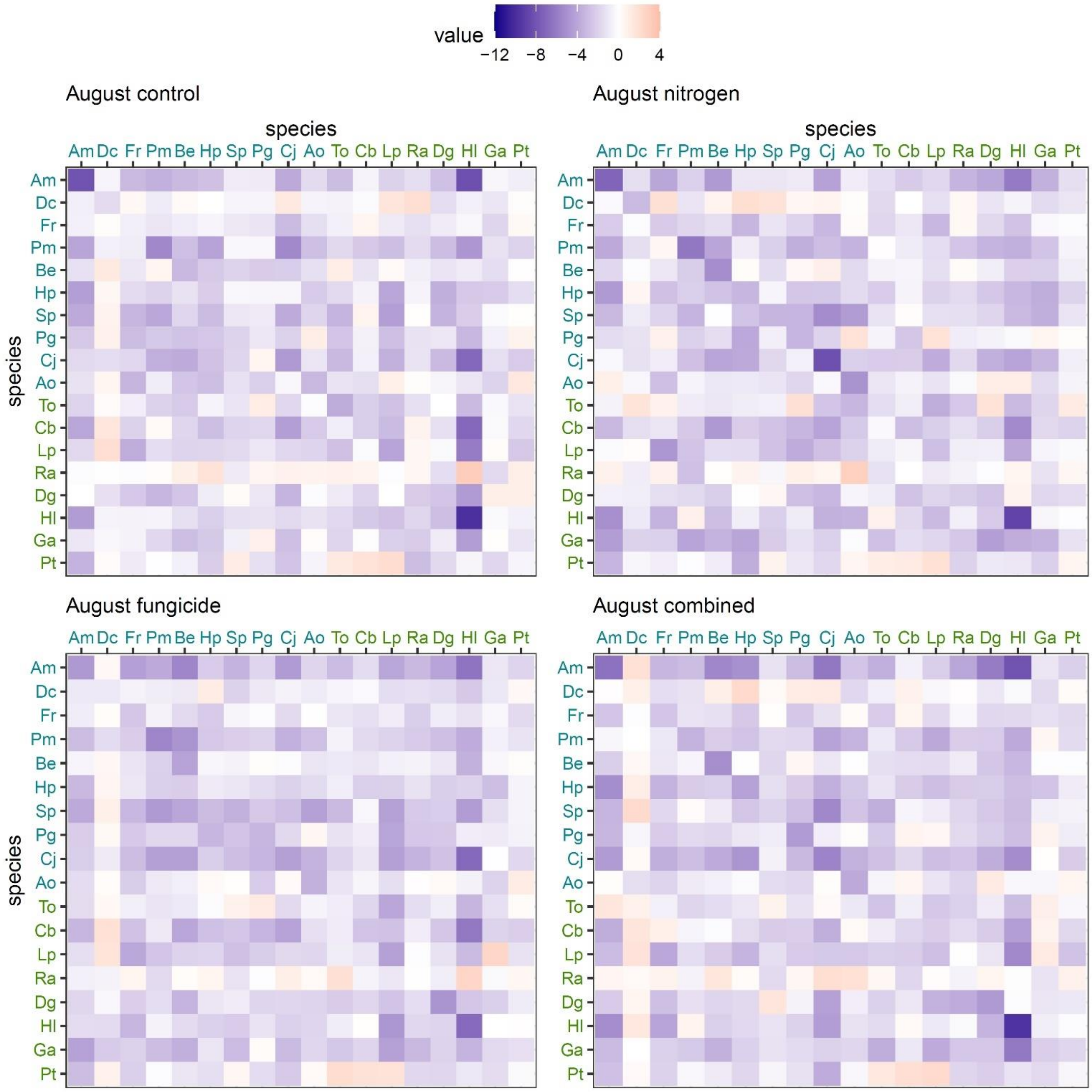


- Figure S1. Matrices of pairwise interactions between the 18 species sampled in our experiment in August. One matrix was obtained for each treatment. Each cell shows a pairwise interaction, colours show the sign and the magnitude of the interaction. Species names in blue are slow growing species, green are fast growing species. Rows show species response to competition from others, columns show species effects on others. See Supplementary Table S1 for species names abbreviation.

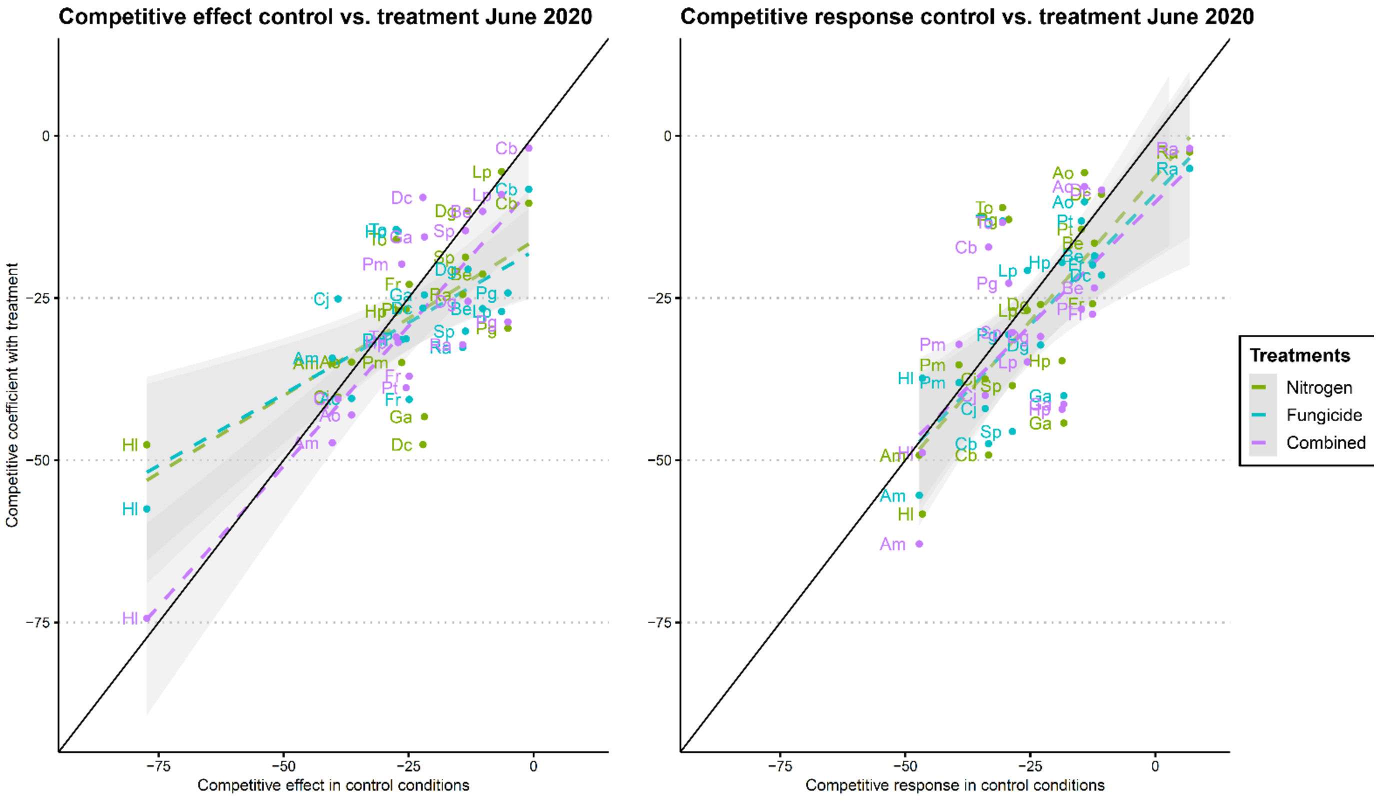


- Figure S2. PaNDiv species competitive effect and response in control versus different treatments (green = nitrogen addition, blue = fungicide addition, purple = combined nitrogen and fungicide addition) in June 2020. Linear regression is represented for each treatment by a dotted line of the corresponding colours (the differences between treatments were not significant). See Supplementary Table S1 for species names abbreviation.

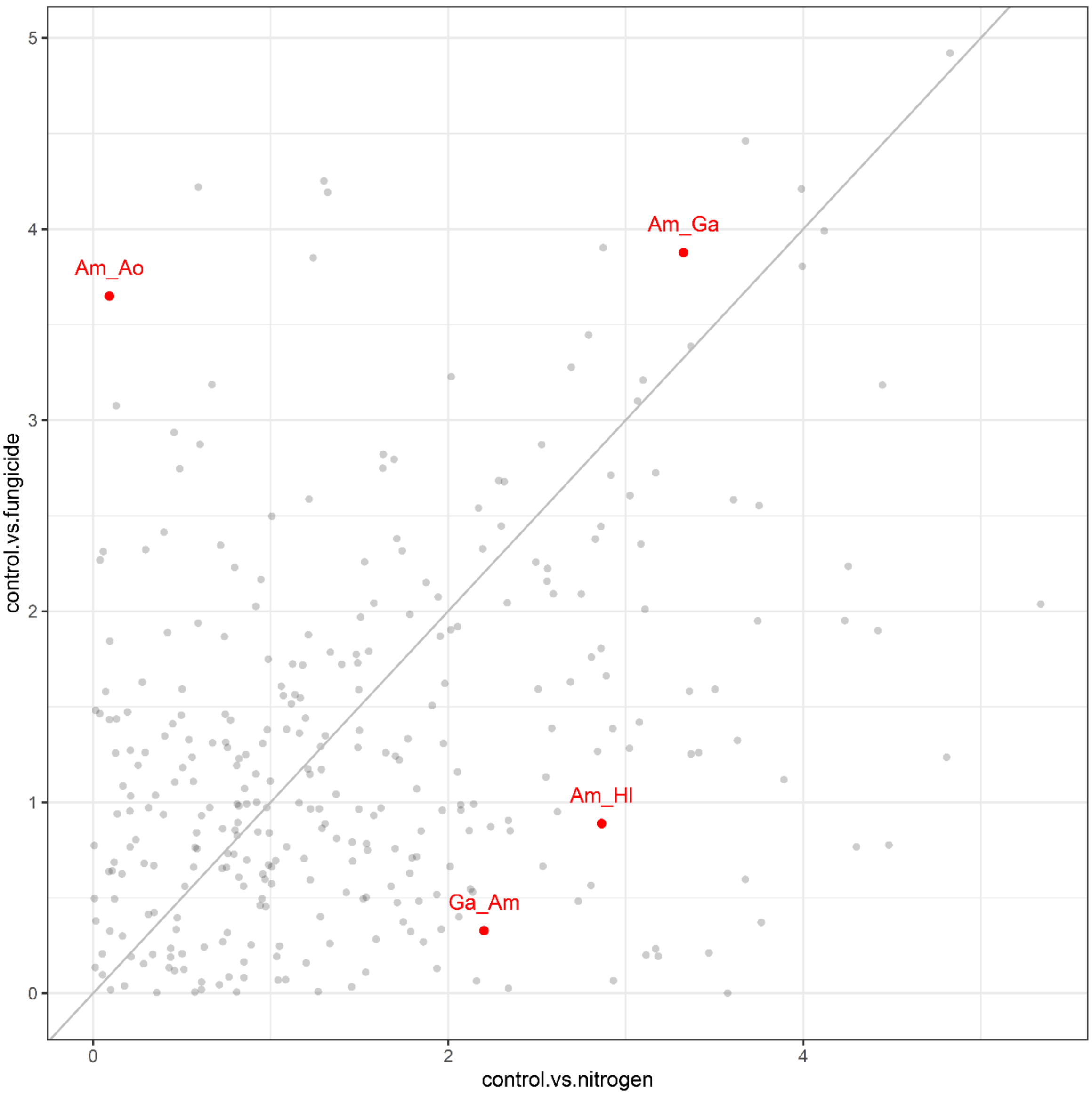


- Figure S3. Difference in pairwise interaction coefficients (in absolute values) between control and nitrogen or control and fungicide treatments. The grey line shows the pattern we would expect if both changes were correlated (no pattern observed here). We highlighted some pairwise interactions in red as examples: notably *Am_Ga* the competitive effect of *Ga* on *Am*, which is modified by both treatment (close to the grey line). In contrast, its reciprocal *Ga_Am* the competitive effect of *Am* on *Ga*, is primarily modified by nitrogen addition. See Supplementary Table S1 for species names abbreviation.

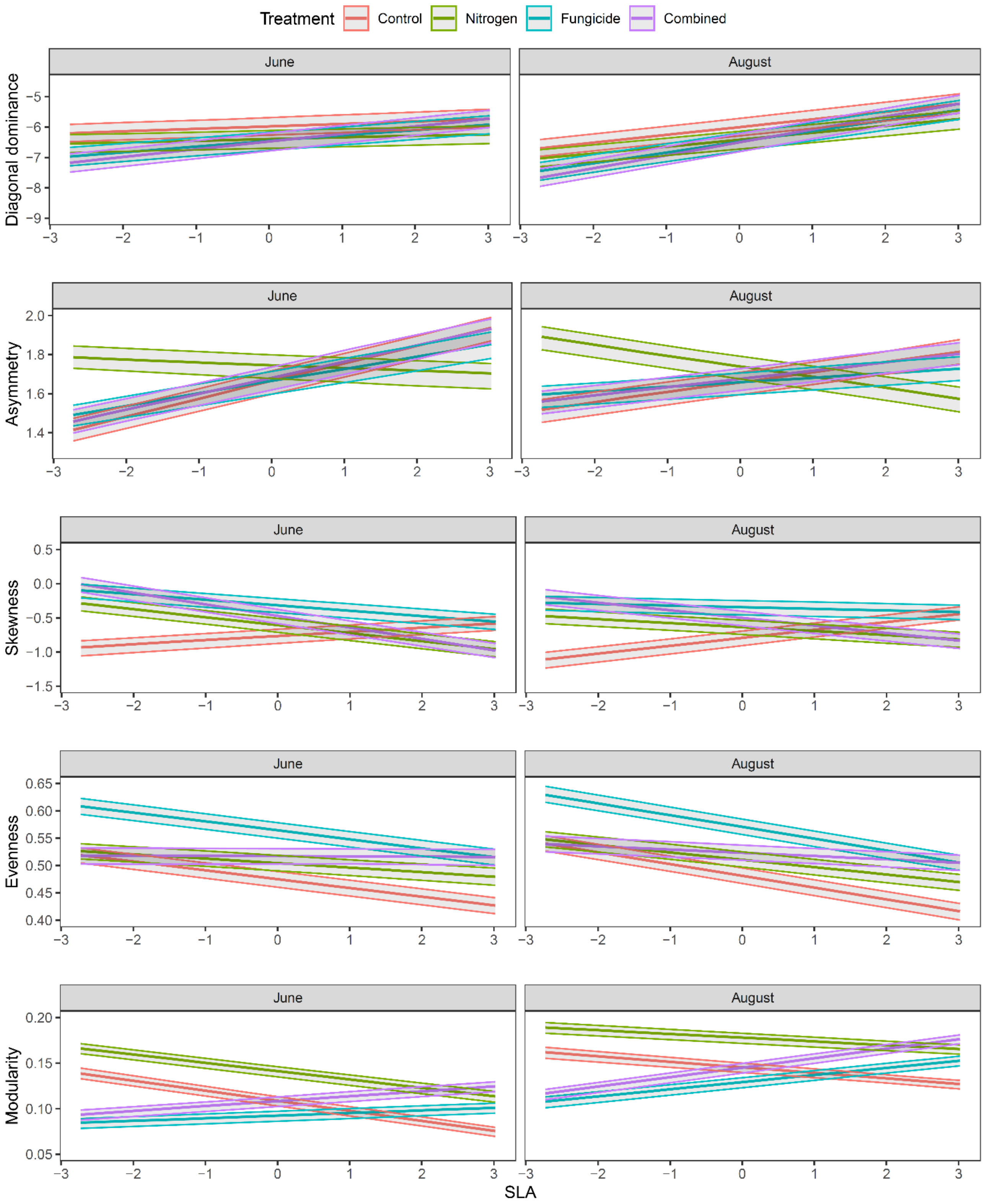


- Figure S4. Impact of SLA and treatments on network metrics for communities of 5 PaNDiv species in June (left) and August (right) 2020. For each network metric, colours determine treatments, and model predictions are shown in the form of predicted mean (bold lines) and confidence intervals (upper and lower lines). SLA was scaled for comparison. SLA = specific leaf area.

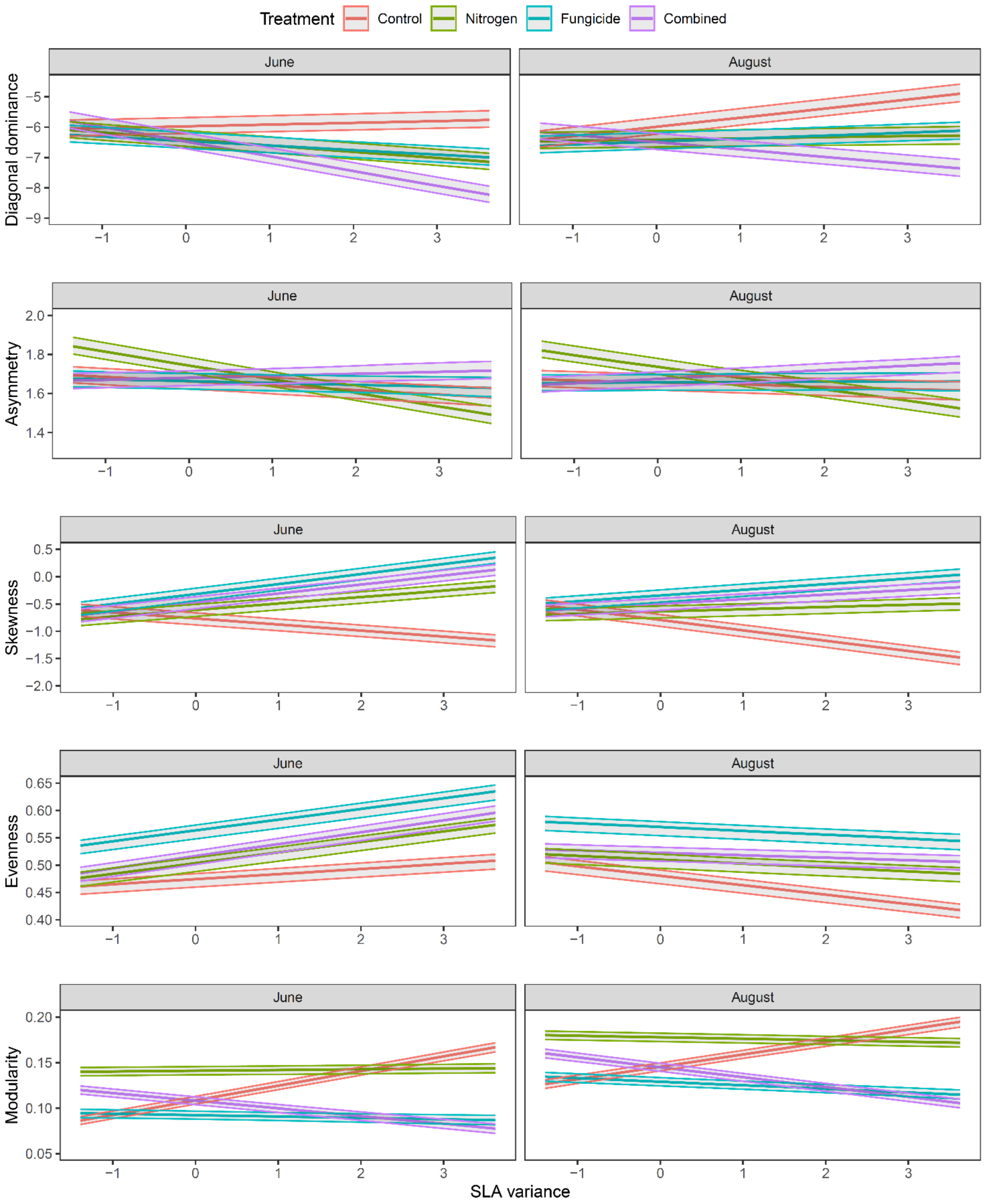


- Figure S5. Impact of SLA variance and treatments on network metrics for communities of 5 PaNDiv species in June (left) and August (right) 2020. For each network metric, colours determine treatments, and model predictions are shown in the form of predicted mean (bold lines) and confidence intervals (upper and lower lines). SLA was scaled for comparison. SLA = specific leaf area.

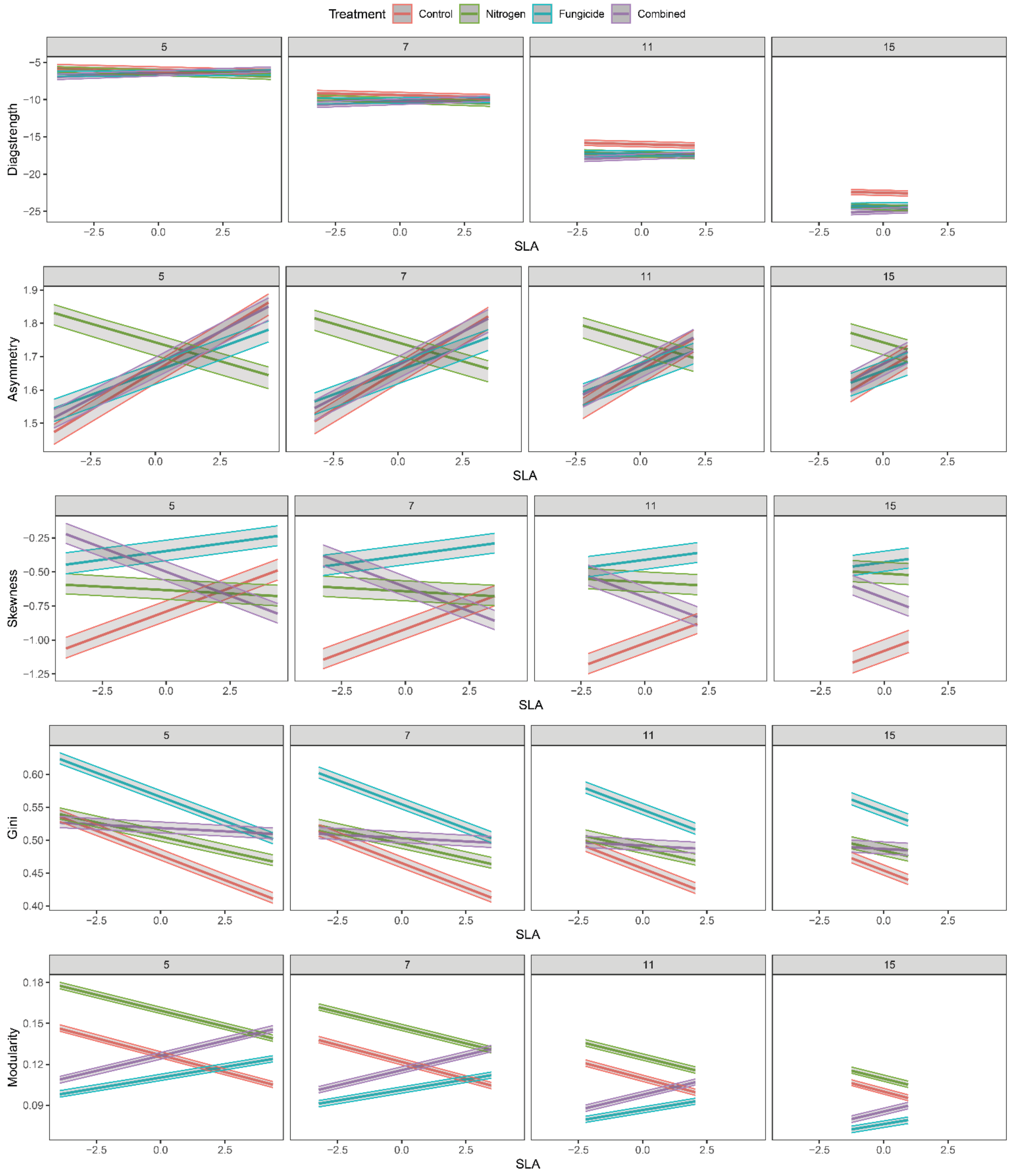


- Figure S6. Evolution of the impact of SLA and treatment on network metrics for networks of different sizes : 5, 7 11 and 15 species in June 2020. For each network metric, colours determine treatments, and model predictions are shown in the form of predicted mean (bold lines) and confidence intervals (upper and lower lines). SLA was scaled for comparison. SLA = specific leaf area.

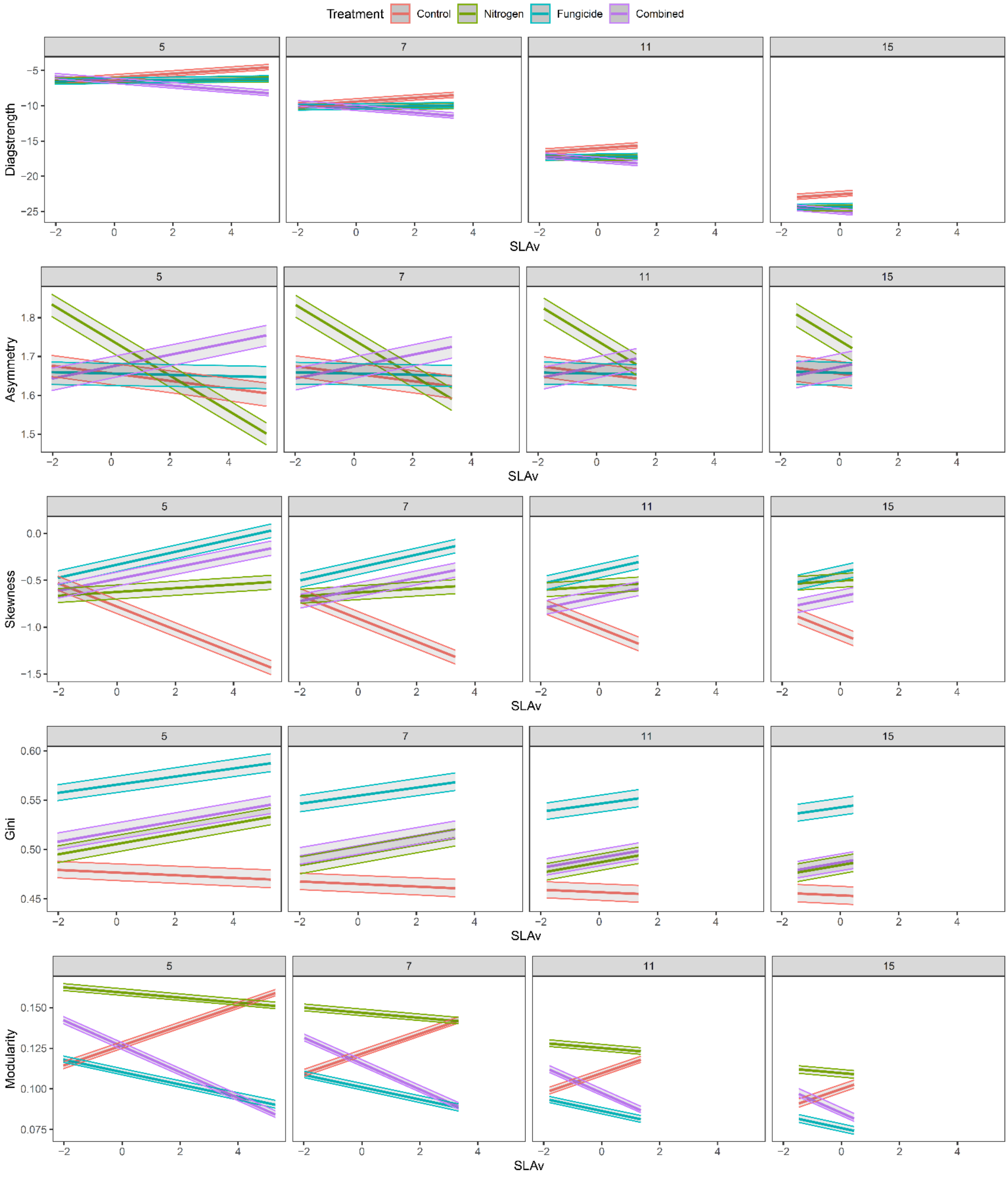


- Figure S7. Evolution of the impact of SLA variance and treatment on network metrics for networks of different sizes : 5, 7 11 and 15 species in June 2020. For each network metric, colours determine treatments, and model predictions are shown in the form of predicted mean (bold lines) and confidence intervals (upper and lower lines). SLA variance was scaled for comparison. SLA = specific leaf area.

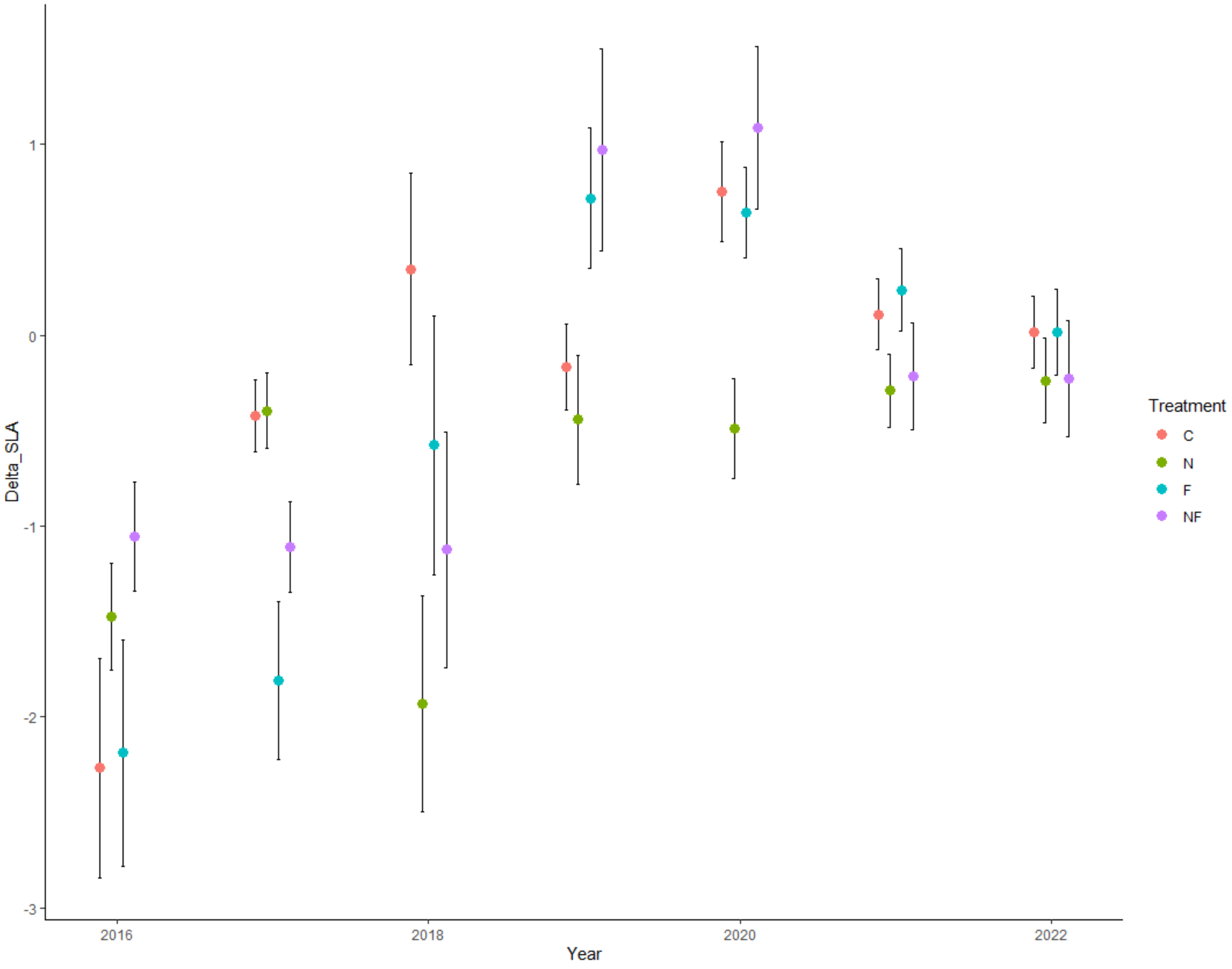


- Figure S8. Shift in SLA of PaNDiv communities in control conditions (orange) with nitrogen addition (green), fungicide addition (blue), and combined addition of both treatments (purple) in June between 2016 and 2022. Negative values of Delta_SLA indicate a shift toward more slow growing communities. SLA = specific leaf area.

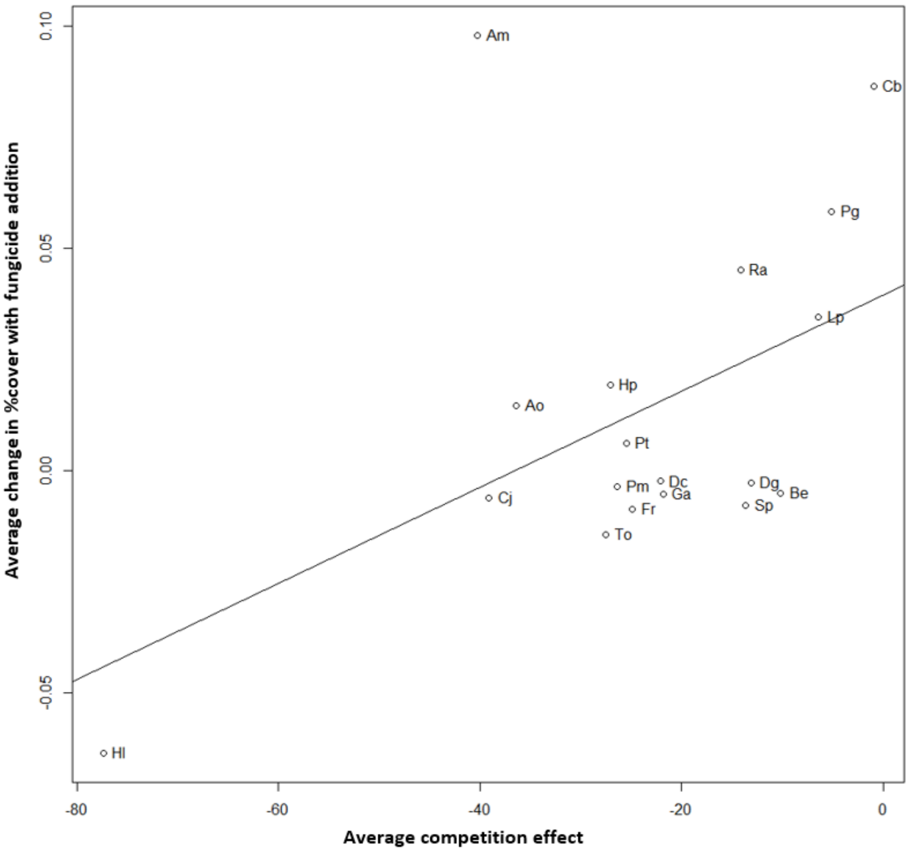


| | diff | | |
|---|---|---|---|
| *Predictors* | *Estimates* | *CI* | *p* |
| (Intercept) | 0.0394 | 0.0093 – 0.0694 | **0.013** |
| competition effect | 0.0011 | 0.0001 – 0.0021 | **0.040** |
| Observations | 18 | | |
| $R^2$ / $R^2$ adjusted | 0.239 / 0.191 | | |

- Figure S9. Linear regression between species increase or decrease in field percentage cover when adding fungicide in PaNDiv experiment (June 2020) and their average competitive effect in control conditions for the same sampling period. See Supplementary Table S1 for species names abbreviation.
